# Metabolomic changes underpinning permethrin resistance in the *Anopheles gambiae* malaria vector from Cameroon

**DOI:** 10.64898/2026.01.13.699336

**Authors:** Steve V. Djova, Michael W. Christopher, Sulaiman S. Ibrahim, Mersimine Kouamo, Rhoel R. Dinglasan, Timothy J. Garrett, Charles S. Wondji

## Abstract

Insecticide resistance reduces the effectiveness of current malaria vector control interventions. To understand the mechanisms underlying permethrin resistance in field-derived *Anopheles gambiae* vectors collected from Cameroon, we used Ultra-High Performance Liquid Chromatography coupled with High-Resolution Tandem Mass Spectrometry (UHPLC-HRMS/MS) to comprehensively analyze metabolic changes in resistant and susceptible samples to gain insight into mechanisms driven permethrin resistance. Resistant mosquitoes exhibited upregulated levels of inosine, nicotinic acid, dipeptides, amino acids, fumarate, uracil, and aldopentose. These data suggest that altered metabolic-based detoxification, as well as target site and metabolic shifts, and enhanced energy production contribute to permethrin resistance. Conversely, susceptible mosquitoes showed increased levels of N-acetyl-aspartic acid, xanthurenic acid, 2-hydroxyglutarate, 3-hydroxykynurenine, propanoylcarnitine, and L-pipecolic acid. These metabolites are associated with neurotoxicity, energy disruption, as well as tryptophan and lysine catabolism. These findings elucidate the metabolic pathways of permethrin-resistance and underscore the mechanisms that could lead to the emergence of pyrethroid cross-resistance.

## Introduction

Malaria remains a significant global health concern, with 597,000 deaths reported in 2023 alone^1^. Sub-Saharan African countries continue to carry the heaviest burden of mortality, with 95% of the estimated malaria deaths worldwide, and 76% of those deaths among children under the age of 5 caused primarily by *Plasmodium falciparum* ^1^. Malaria prevention still relies heavily on the use of insecticide treated nets (ITNs) and indoor residual spraying (IRS) for vector control. However, the effectiveness of these tools is currently challenged by the spread of pyrethroid resistance across major malaria mosquito vectors, such as *Anopheles gambiae*^2,3^ and *Anopheles funestus*^4–6^. Mosquitoes have evolved a variety of ways to detoxify pyrethroid insecticides, with the two main mechanisms being (i) target site resistance, known as knockdown resistance (*kdr*), a mutation in the voltage-gated sodium channel that reduces pyrethroid binding and subsequent toxicity in mosquitoes^7–9^; and (ii) metabolic resistance through increased gene expression/amplification or allelic variation of cytochrome P450s, carboxylesterases (COEs), glutathione S-transferases (GSTs) and other oxidation/reduction genes^10,11^. While extensive ITN distribution and coverage are primary drivers of pyrethroid resistance in Cameroon^2,12^, evidence suggests the involvement of other contributing factors. Notably, the application of pesticides in agriculture may increase pyrethroid tolerance in mosquitoes^13^. A case in point is Nkolondom, a peri-urban agricultural site in Yaoundé, Cameroon, where routine pesticide application by farmers has increased pyrethroid resistance in mosquitoes^14^.

Progress in understanding pyrethroid resistance mechanisms in *An. gambiae sensu lato (s.l.*) has revealed both target-site resistance, through the *kdr* (L1014F) alleles^2,14,15^, and metabolic resistance, associated with the upregulation of detoxification genes including P450s (*cyp6P3*, *cyp6p4*, *cyp6z3*, *cyp9k1*) and glutathione S-transferases genes (*gste2*, *gsttd1-6*, *gstd1-4*)^16–19^. However, the metabolic changes accompanying the up regulation of these resistance genes and important biomarkers of resistance (such as key metabolites) are still unknown. *In vitro* metabolism assays with recombinant enzymes have confirmed the role of many cytochrome P450 enzymes in pyrethroid resistance in *An. gambiae* ^20^. Stevenson *et al.*^21^, used recombinant *An*. *gambiae* cytochrome P450 CYP6M2 to perform an *in vitro* analysis of deltamethrin metabolism. Their finding revealed a sequential breakdown of deltamethrin with 4’-hydroxylation identified as a major metabolism route. The study of Ouafo *et al.*^22^ found that a recombinant Cyp6p3-E205D from *An. gambiae* has a 2.5-fold increase in its ability to metabolize alpha-cypermethrin and permethrin insecticides. While *in vitro* functional assays make identifying primary pyrethroid metabolites straightforward, the detoxification process in mosquitoes is more complex. Previous research have identified new features of pyrethroid detoxification in mosquito vectors^23^. Permethrin resistance in *Drosophila melanogaster* involves insecticide detoxification and multiple cellular pathways, with metabolomic alterations in tryptophan catabolism as a secondary target^24^. Building on these findings, we used UHPLC-HRMS/MS to conduct untargeted metabolite profiling to measure metabolome-associated changes in permethrin-resistant field-derived *An. gambiae sensu stricto (s.*s.) from Cameroon.

## Results

### Metabolic response of *An. gambiae s.s.* from Nkolondom exposed to permethrin 0.75%

All mosquitoes from Nkolondom were confirmed by PCR as *An. gambiae s.s.* and WHO bioassays test tubes revealed permethrin resistance for these mosquitoes with 76.3% mortality after 1h, and 25.8% mortality after 30 min (0.5h) exposure to permethrin 0.75% **(Figure 1A)**. Our untargeted analysis identified a large number of features and adducts in positive and negative ionization **(Tables S1 and S2)**. From ANOVA, we found 2,767 significant features including adducts in negative ion mode and 2,524 significant features in the positive ionization mode **(Tables S3 and S4, respectively**). Principal component analysis (PCA) indicated that the metabolome resulted in three clusters, alive at 0.5h, dead at 0.5h, and unexposed clustering together; alive at 1h, and dead at 1h clusters **(Figure 1B)**. Sparse partial least squares-discriminant analysis (sPLS-DA), (performance showed in **Figure S1)**, successfully resolved four distinct clusters corresponding to mosquitoes that were alive at 1h, dead at 1h, dead at 0.5h, and a combined group of unexposed (controls) and alive at 0.5h mosquitoes **(Figure 1C)**. These sPLS-DA demonstrate the distinct metabolic responses associated with permethrin (0.75%) exposure over time.

**Figure 1:**
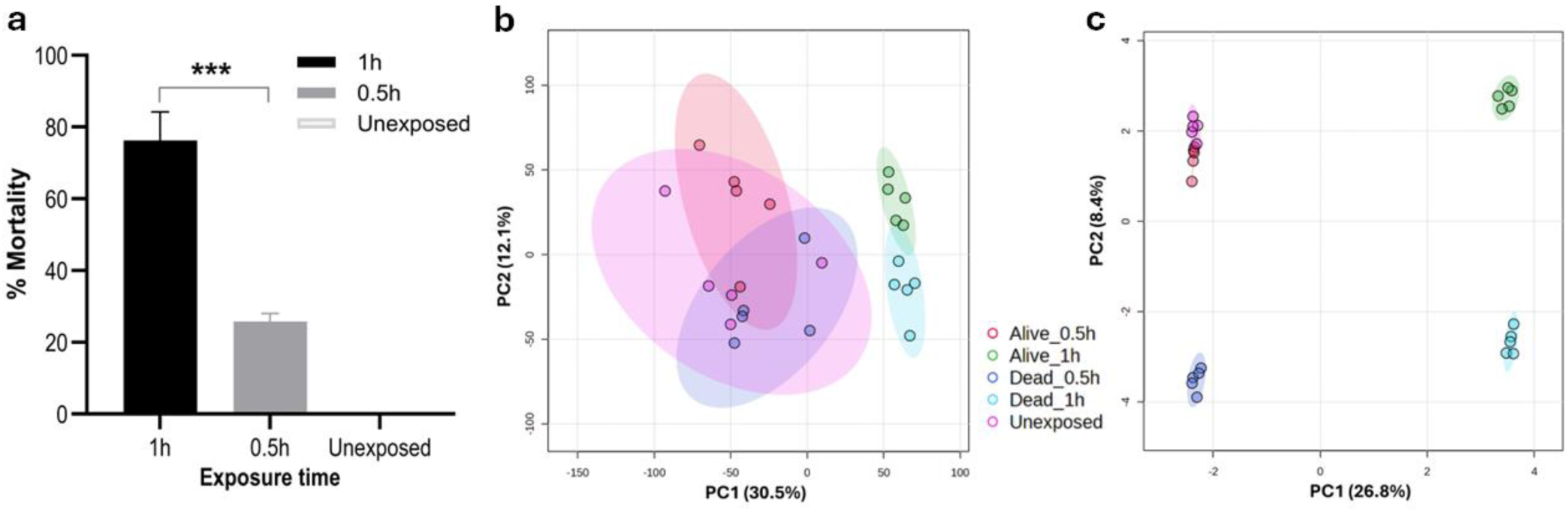
Metabolic response of *A. gambiae* s.s. to permethrin exposure. (a) Susceptibility profile of *An. gambiae s.s.* collected in Nkolondom and exposed during 1h and 0.5h to permethrin 0.75%. The data shown are mean ± SD, N=100 at 1h and 0.5h, ANOVA, post-hoc Fisher’s tests ***p˂0.0001. (b) Principal Component Analysis (PCA) of the metabolome resulted in three clusters from (unexposed, alive at 0.5h, dead at 0.5h clustered together; alive at 1h, and dead at 1h). Sparse Partial Least Squares-Discriminant Analysis (sPLS-DA) provided greater discrimination, identifying four distinct clusters, unexposed and alive at 0.5h clustered together, dead at 0.5h; dead at 1h, and alive at 1h clusters.

UHPLC-HRMS/MS analysis revealed distinct metabolite profiles when comparing permethrin-resistant, to susceptible mosquitoes. In the first column cluster, resistant and unexposed mosquitoes exhibited elevated levels of L-phenylalanine, L-isoleucine, L-methionine, and the dipeptides alanyl-L-methionine and glycyl-L-methionine while the level was low in susceptible phenotypes. Conversely, in the second column cluster, dead, susceptible mosquitoes showed increased levels of xanthurenic acid (XA), 2-hydroxyglutarate, propanoylcarnitine, L-pipecolic acid, N-acetyl-aspartic acid (NAA), hexanoylcarnitine, N-acetyl-glutamate, N-acetyltyramine, butanoylcarnitine, acetyl-L-carnitine, 3-hyhydroxykynurenine, and orotate while their concentration was low in resistant and unexposed phenotypes **(Figure 2a)**. Negative ion mode also confirmed in the first column cluster, higher levels of 5-hydroxy-L-tryptophan, fumarate, aldopentose, nicotinate, inosine, L-threonine, L-tryptophan, L-methionine, L-isoleucine, L-valine, and uracil in resistant and unexposed mosquitoes. In the second column cluster, N-acetyl-aspartate, mannitol, xanthurenic acid, orotate and 2-methylglutarate revealed elevated level in dead susceptible mosquitoes compared to resistant and unexposed groups **(Figure 2b)**.

**Figure 2:**
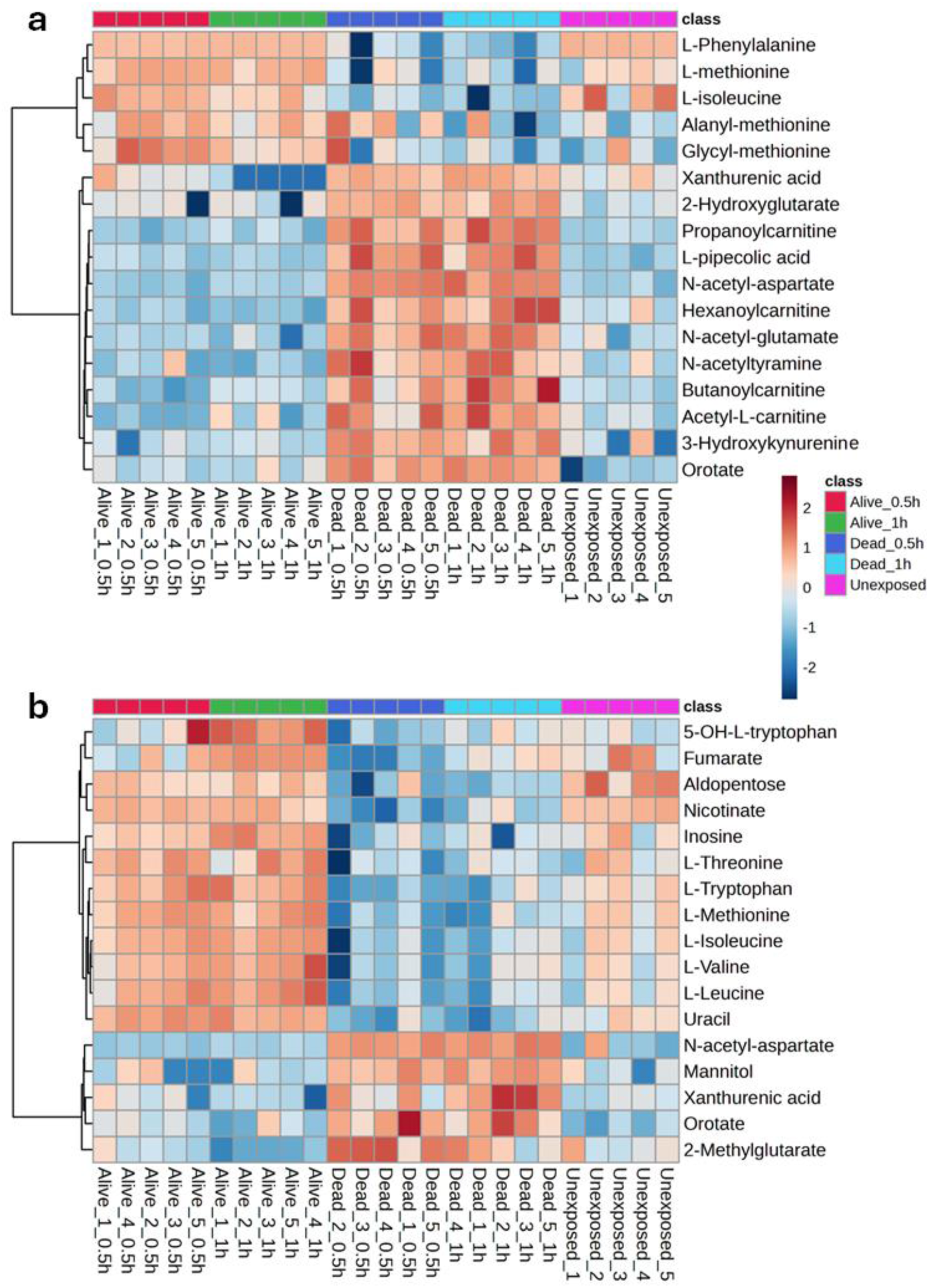
Heatmaps illustrate the most significant differences in the mosquito metabolomic profile across five biological replicates for each phenotype. Positive mode (A) and Negative mode (B).

### Metabolite markers of permethrin detoxification in permethrin-resistant *An. gambiae*

While metabolome profiling for permethrin detoxification presents inherent challenges, variations in critical metabolite abundance provide valuable insights into the specific enzymatic pathways involved. Notably, resistant mosquitoes exhibited elevated levels of nicotinate **(Figure 3a, Figure S2)**, a precursor for NAD+ and NADP coenzymes, both indispensable for cytochrome P450 activation, regeneration, and metabolic energy generation. We also observed the higher levels of uracil **(Figure 3b, Figure S3)** in resistant mosquitoes. Additionally, the upregulation of inosine in resistant mosquitoes, as revealed in **Figure 3d, (Figure S4)**. Conversely, susceptible mosquitoes exhibited elevated orotate levels **(Figure 3c, Figure S5)**

**Figure 3:**
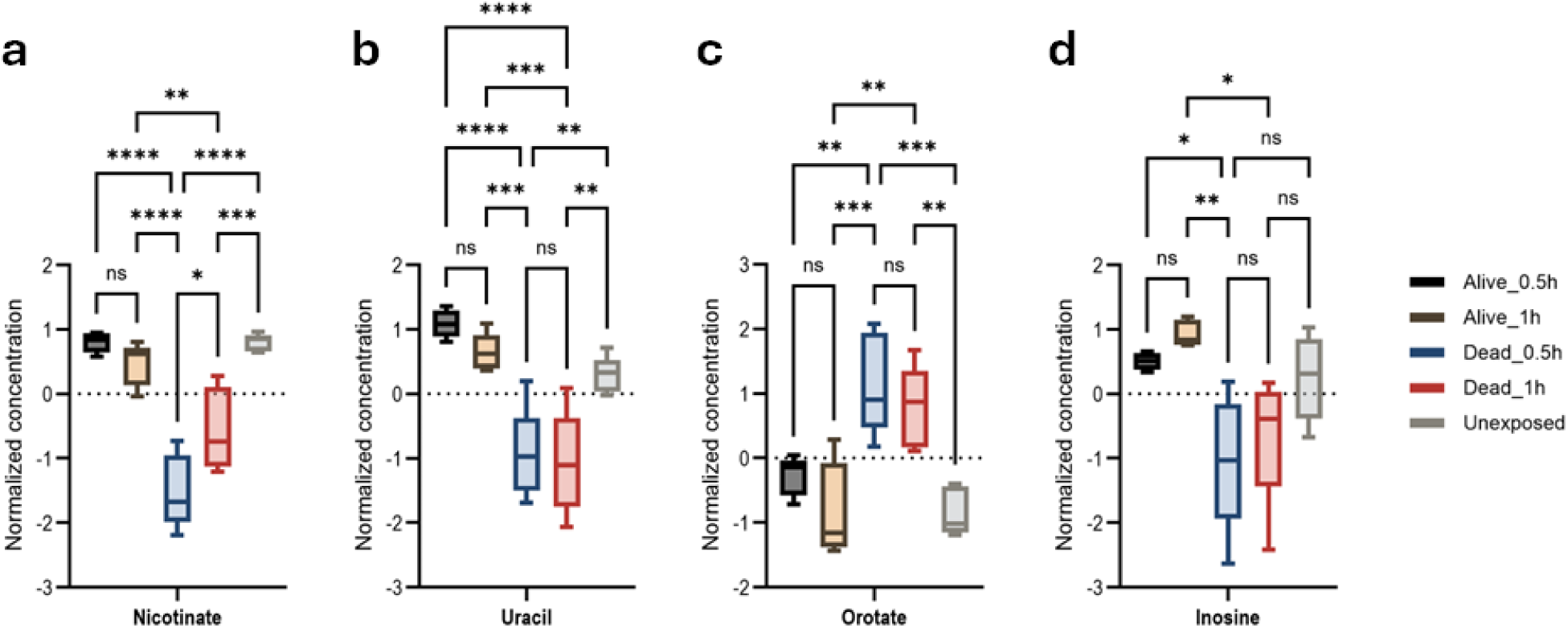
Confirmation of metabolic and target site resistance, as well as purine and pyrimidine metabolism pathways associated with *An. gambiae* permethrin-resistance phenotypes. Anova, post-hoc Fisher with p-value threshold=0.05, statistical annotation: ns, non-significant. (a) Nicotinate *p-value=0.0146; **p-value=0.0082; ***p-value=0.0005; ****p-value ˂0.0001. (b) Uracil **p-value=0.0094; ***p-value=0.0008, ****p-value ˂0.0001. (c) Orotate **p-value=0.0029; ***p-value=0.0004. (d) Inosine *p-value=0.0131; **p-value=0.0018.

### Amino acid profiles differentiate *An. gambiae* permethrin-resistance phenotypes

This study associated elevated levels of the dipeptides Alanyl-L-methionine and Glycyl-L-methionine, along with the free amino acids L-methionine, L-isoleucine, L-leucine, L-phenylalanine, L-tryptophan, L-valine, and L-threonine, with permethrin-resistance in *An. gambiae* mosquitoes **(Figure S6-15)**.

### Energy metabolism is swift in resistant phenotypes

Several key metabolites that are involved with energy reserve catabolism were identified. Resistant and unexposed mosquitoes maintained higher aldopentose and fumarate levels than susceptible mosquitoes, indicating unaffected energy production. In contrast, all mosquitoes that died following 0.5h or 1h exposure, showed significantly elevated mannitol **(Figure 4a, b, c, Figure S16-18)**.

**Figure 4:**
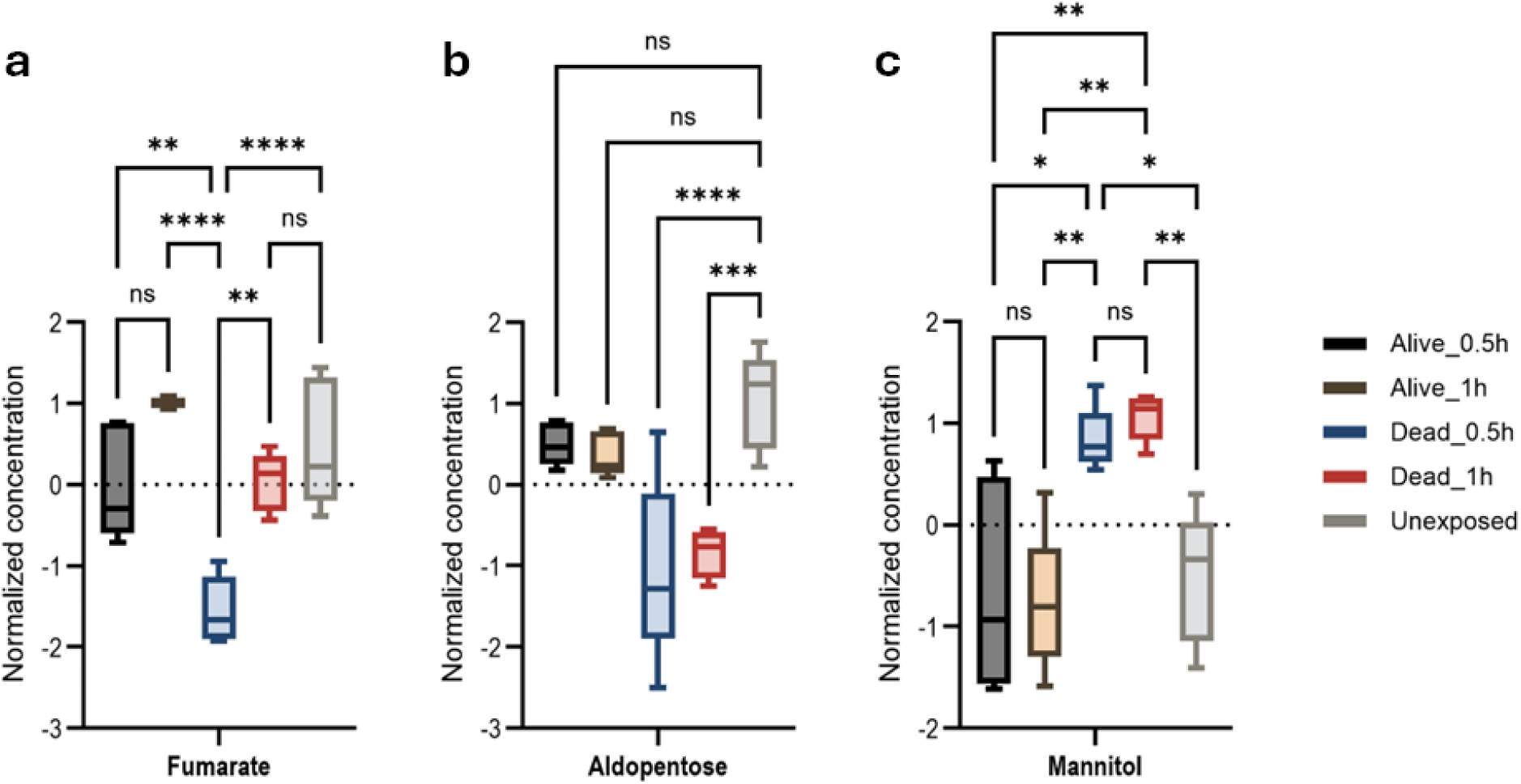
Energy metabolism responses associated with *An. gambiae* permethrin-resistance phenotypes. Anova, post-hoc Fisher with p-value threshold=0.05, statistical annotation: ns, non-significant. (a) Fumarate, **p=0.0015, ****p-value˂0.0001; (b) Aldopentose, ***p-value=0.0003, (c) Mannitol *p-value=0.0154, **p-value=0.0021.

### Hypoxia markers associated with permethrin neurotoxic effects

L-2-hydroxyglutaric acid and 2-methylglutaric acid were significantly elevated in susceptible mosquitoes compared to resistant and unexposed groups, as shown in **Figure S19-21**.

### Amino acid N-acetylation pathway associated with susceptible phenotypes

N-acetyl-L-glutamic acid, N-acetyl-tyramine, and N-acetyl-L-aspartic acid (NAA) were significantly elevated in susceptible mosquitoes **(Figure 5a, b, c, and Figure S22-24)**. We used an NAA standard to confirm the chromatographic retention times of 2.09 min and MS/MS spectra to validate its exclusive presence among dead mosquito samples **(Figure S25)**.

**Figure 5:**
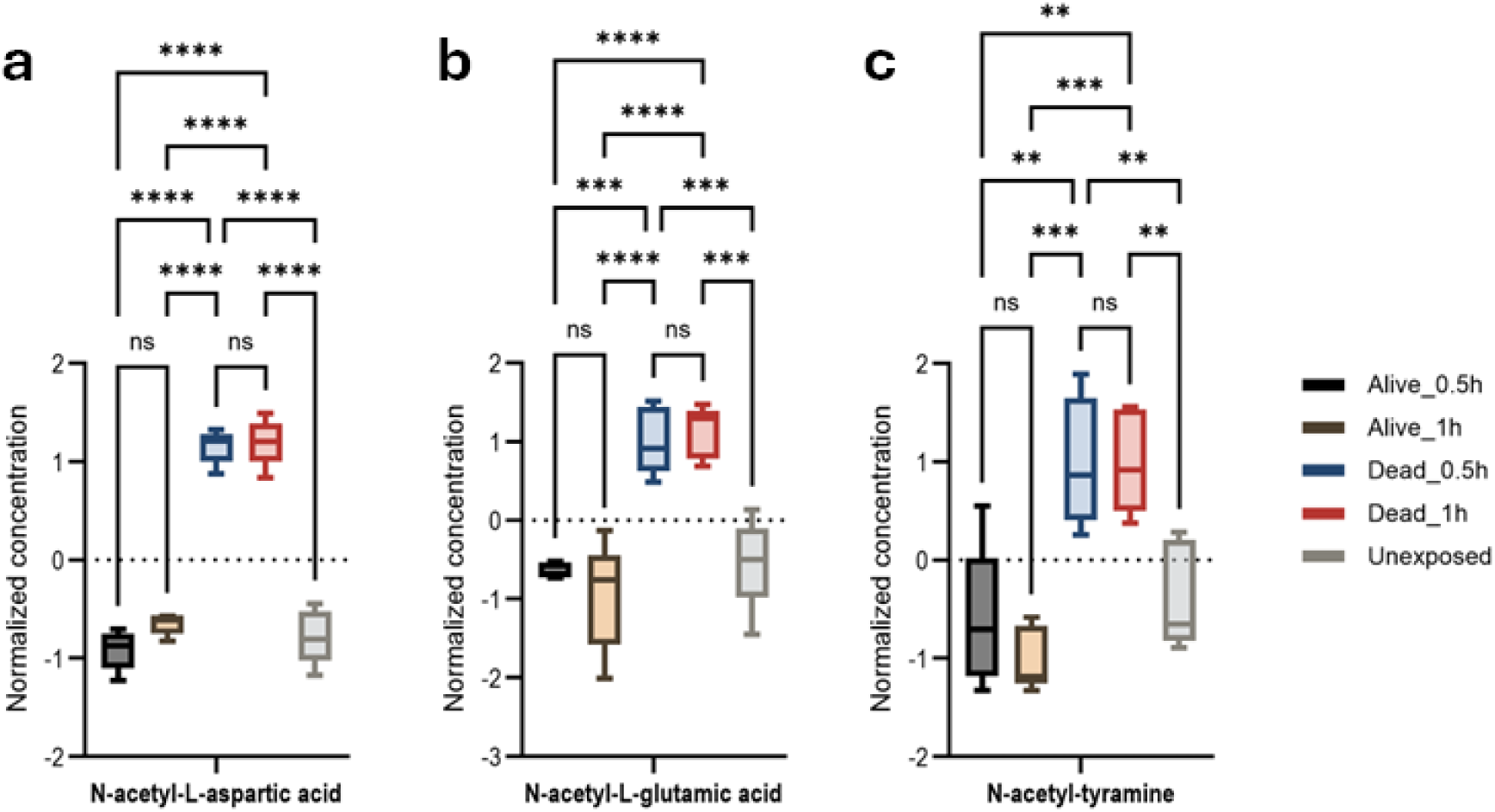
Protein N-acetylation and degradation associated with *An. gambiae* permethrin-resistance phenotypes. Anova, post-hoc Fisher p-value threshold=0.05, statistics annotation: ns, non-significant, (a) N-acetyl-L-aspartic acid, ****p-value ˂0.0001; (b) N-acetyl-L-glutamic acid, ***p-value=0.0002, ****p-value ˂0.0001; (c) N-acetyl-tyramine, **p-value=0.002, **p-value=0.0002.

### Tryptophan and lysine metabolism pathways associated with susceptible phenotypes

Levels of xanthurenic acid (XA) and 3-hydroxykynurenine, neurotoxic biomarkers and tryptophan catabolites (**Figure 6a, b, Figure S26-27**), were elevated in susceptible mosquitoes compared to those resistant at 0.5h and 1h). The upregulation of 5-hydroxy-L-tryptophan, precursor of serotonin neurotransmitter in resistant 1h group may indicate a potential action in permethrin resistance (**Figure 6c, Figure S28**. L-pipecolic acid (L-PA), catabolite of lysine, was also elevated in the susceptible group **Figure 6d, S29**).

**Figure 6:**
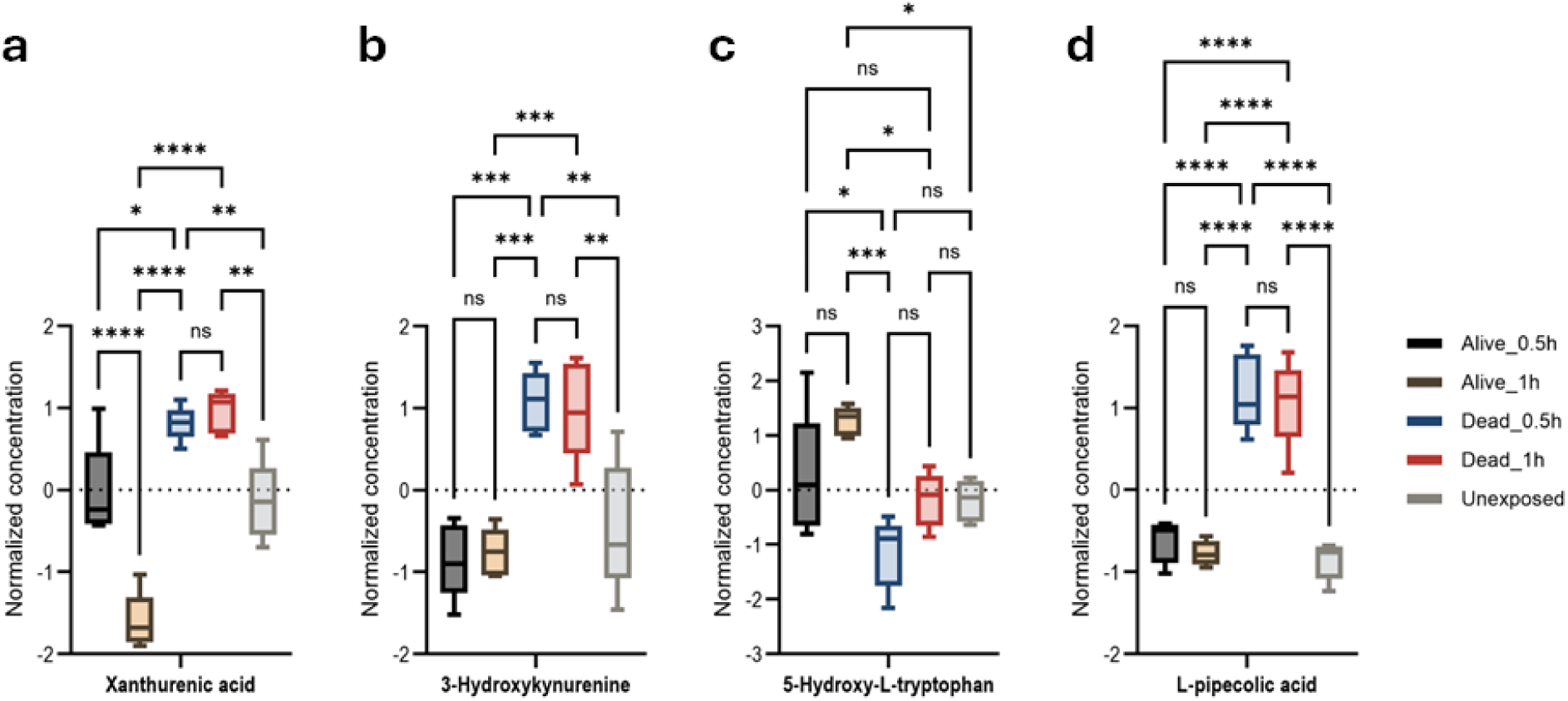
Tryptophan and lysine catabolism pathway associated with permethrin *An. gambiae* permethrin-resistance phenotypes. Anova, post-hoc Fisher p-value threshold=0.05, statistics annotation: ns, non-significant, (a) XA, *p-value=0.0257, **p-value=0.0072, ***p-value=0.0001, ****p-value ˂0.0001; (b) 3-hydroxykynurenine **p-value=0.0018, ***p-value=0.0001; 5-(c) hydroxy-L-tryptophan, *p-value=0.018, ***p-value=0.0001; (d) L-PA, ****p-value ˂0.0001.

### Carnitine/acylcarnitine shuttle alteration

A mosquito’s capacity to survive permethrin exposure is directly linked to its ability to generate supplemental energy. The findings showed increased levels of hexanoylcarnitine, propanoylcarnitine, butanoylcarnitine, and N-acetyl-L-carnitine exclusively in susceptible mosquitoes (compared to resistant and unexposed groups). This pattern suggests energy depletion and a subsequent alteration of the acylcarnitine shuttle, leading to the accumulation of these short-chain fatty acids in susceptible mosquitoes **(Figure 7a, b, c, d, Figure S30-33)**.

**Figure 7:**
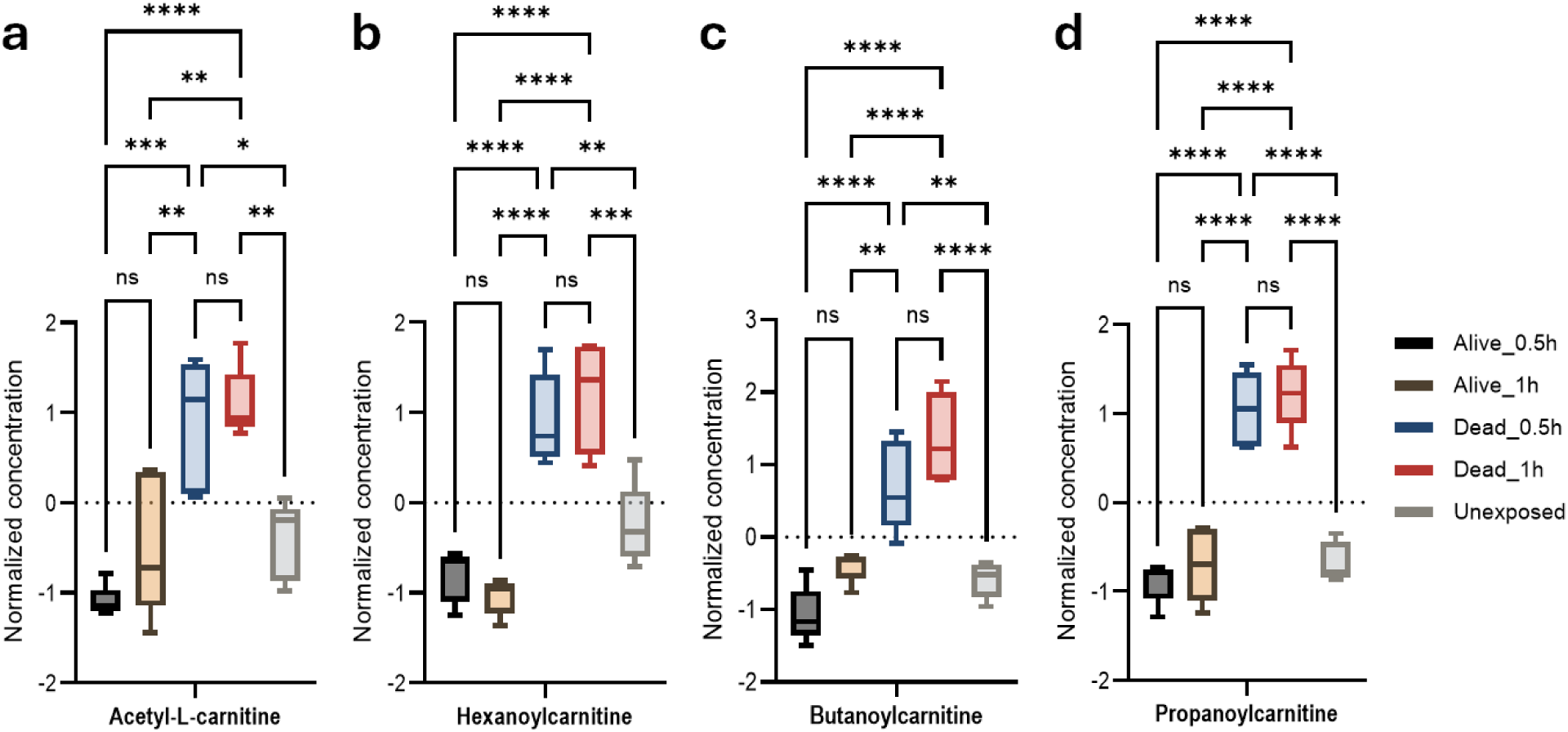
Carnitine/acylcarnitine shuttle associated with permethrin *An. gambiae* permethrin phenotype. Anova, post-hoc Fisher p-value threshold=0.05, statistics annotation: ns, non-significant, (a) Acetyl-L-carnitine, *p-value=0.0113, **p-value=0.0028, ***p-value=0.0001, ****p-value ˂0.0001; (b) Hexanoylcarnitine, **p=0.0034, ***p=0.0004, ****p-value˂0.0001; (c)Butanoylcarnitine, **p=0.0019, ****p-value ˂0.0001; (d) Propanoylcarnitine ****p-value˂0.0001.

## Discussion

Permethrin metabolism is a known multi-step process. Carboxylesterases and cytochromes sequentially act on permethrin, producing intermediates such as 3-phenoxybenzyl alcohol. cytochrome orthologs then convert this alcohol to 3-phenoxybenzaldehyde, followed by aldehyde dehydrogenase transformation into a non-toxic phenoxybenzoic acid ^25–27^. These non-toxic metabolites are subsequently excreted from cells and tissues via efflux pumps ^28^ or the mosquito’s Malpighian tubules^29^. In this study, none of the multiple intermediates of permethrin have been identified as a significant metabolite in a resistant phenotype. This result may reflect the role of ABCG transporters predicted to act as efflux pumps that clear pyrethroids or toxic intermediates from cells or tissue in *An. gambiae* mosquitoes ^30^. Nevertheless, the presence of metabolite signals like cis-3-(2,2-dichlorovinyl)-2,2-dimethyl-(1-cyclopropane) carboxylic acid (*m/z* 206.9926) and phenoxybenzoic acid (3-PBA) (*m/z* 213.0557) indicates permethrin detoxification is likely mediated by carboxylesterases and monooxygenase cytochrome P450 enzymes.

High nicotinic acid levels in resistant mosquitoes, a precursor to NAD+ and NADP+, support the role of monooxygenase cytochrome P450 enzymes in permethrin detoxification, consistent with findings from Hannemann *et al.*^31^ on NAD(P)H-dependent ferredoxin and the fact that monooxygenases are not electron-transfer proteins. The increased nicotinamide levels observed by Brinser *et al.*^24^ in *Drosophila melanogaster* exposed to permethrin suggests that is a conserved metabolic response in Diptera. The elevated uracil levels detected in permethrin-resistant and unexposed mosquitoes, in contrast to susceptible groups, likely fuel RNA synthesis, a process that correlates with P450-mediated permethrin detoxification and could explain the necessity for high uracil levels to support the increased production of detoxifying enzymes ^20^. The accumulation of orotate, an intermediate of UMP synthesis, in susceptible mosquitoes following permethrin exposure suggests a potential disruption of RNA synthesis. This observation aligns with previous research demonstrating permethrin’s impact on pyrimidine metabolism^24^.

Inosine is an intermediate in purine metabolism. Increased inosine levels in resistant mosquitoes may suggest IMP metabolism disruption, xanthine metabolism inhibition or post-transcriptional modification (A-to-I editing) of the pre-mRNA encoding the voltage-gated sodium channel. While two datasets dispute the genotype-phenotype link^32^, our findings align with previous studies showing that A-to-I editing of *para* transcripts in *An. gambiae, Aedes aegypti styl* and *Drosophila* affects invariant amino acids to varying degrees in mRNA thereby, modulating the sodium channel function^33^. This A-to-I editing of *kdr* could explain resistance in mosquitoes and its connection to the genotype-resistance phenotype observed in *Musca domestica*, *Blattella germanica*, *Culex quinquefasciatus*, and *Aedes albopictus*^34,35^.

A significant change in energy metabolism was observed in resistant, unexposed mosquitoes, evidenced by increased aldopentose levels compared to susceptible mosquitoes. This finding suggests the mobilization and utilization of stored glycogen, consistent with the depletion of glycogen energy reserves previously reported upon permethrin exposure^24^.

Permethrin exposed to mosquitoes may disrupt the Krebs cycle, as the observed upregulation of fumarate, a key intermediate in both resistant and unexposed mosquitoes compared to susceptible ones, strongly suggests a perturbation in energy metabolism.

Multiple complex resistance mechanisms result in greater energy consumption and are upregulated when mosquitoes are exposed to permethrin. Stress-induced protein hydrolysis and release of free amino acids is one source of energy production through the tricarboxylic acid cycle in insects^24,36–38^. Previous studies using targeted and untargeted GC/MS metabolomic approaches have shown increased levels of free amino acids in *An. gambiae* mosquitoes and specific to pyrethroids insecticides^24,39^. Compared to unexposed and susceptible mosquitoes, resistant mosquitoes showed increased levels of dipeptides and free amino acids. These elevated amino acid levels may indicate upregulated pathways associated with insecticide resistance, as previously demonstrated by Forman *et al.*^40^. Altered amino acid profiles, including these changes, are established markers of resistance ^36,41^.

In mosquitoes, the carnitine/acylcarnitine shuttle, facilitating the transport of short-chain fatty acids from the cytoplasm into the mitochondrial matrix for beta-oxidation and ATP production. ^36,42^. Similarly, acylcarnitine is a major intermediate in *Aedes aegypti*, mediating fatty acid-CoA transport for mitochondrial beta-oxidation and ATP generation^42^.

In permethrin-resistant *An. gambiae*, increased energy demands during detoxification may drive elevated fatty acid influx via the carnitine/acylcarnitine shuttle. Notably, susceptible mosquitoes exhibited altered carnitine/acylcarnitine shuttle activity and elevated levels of hexanoylcarnitine, propanoylcarnitine, butanoylcarnitine and N-acetyl-carnitine compared to resistant and unexposed mosquitoes. The observed carnitine/acylcarnitine shuttle alterations point to potential energy production disruptions and a possible role in modulating permethrin resistance in *An. gambiae*.

One of the key findings was the upregulation of XA and the 3-hydroxykynurenine, a bypass product of tryptophan in susceptible mosquitoes compared to resistant and unexposed groups. Xanthurenic acid is also a byproduct of the ommochrome pathway involved in mosquito eye pigmentation and is associated as a signaling molecule to Plasmodium gametocytes to undergo gametogenesis inside the mosquito midgut ^43^. Brinzer *et al.*^24^ attested that permethrin susceptibility in *Drosophila melanogaster* is mediated by the production of XA. Furthermore, they found that the knockdown of the CG6950 gene, which is involved in tryptophan catabolism, increased resistance to permethrin. Our findings confirm that the production of XA and a neurotoxic 3-hydroxykynurenine through tryptophan catabolism makes *Anopheles gambiae* more sensitive to permethrin.

We found a high concentration of NAA, N-acetyl-L-glutamic acid, and N-acetyl-tyramine in susceptible mosquitoes. NAA is present in flies and plays a crucial role in cellular energy production, particularly during stress or nutrient deprivation, by influencing the overall brain energy state in the Drosophila nervous system^44^. This finding suggests a potential role of NAA in disrupting the state of brain energy in *An. gambiae* when exposed to permethrin.

On the other hand, NAA is a precursor of N-acetyl-L-aspartylglutamate (NAAG), a prominent peptide neurotransmitter in the mammalian nervous system^45^. While NAAG has been detected in the *Drosophila* nervous system, with concentrations depending on the fly’s condition^44,46^, its presence and function in the *An. gambiae* nervous system remain unknown. Although NAAG itself has not been detected in *Drosophila*, the related tripeptide NAAG2 has been identified using LC/MS by Kozik^47^. Furthermore, enzymes capable of N-acetylation reactions are present in *Drosophila* ^48,49^. This finding warrants further investigation into the potential role of NAA in modulating permethrin sensitivity in this key malaria vector.

Permethrin’s neurotoxic effects, inducing muscle spasms and paralysis ^9,50^, likely may compromise respiratory function, leading to hypoxia. This hypoxia may correlate with the observed L-2-hydroxyglutarate and 2-methylglutarate accumulation in susceptible mosquitoes, a metabolites known to elevate under hypoxic stress and contribute to mitochondrial dysfunction and lethality in *Drosophila melanogaster* ^51^.

L-PA, a cyclic amino acid derived from lysine catabolism and known to act as an Osmo-protectant in *Corynebacterium glutamicum* ^52^, was upregulated in susceptible mosquitoes compared to resistant and unexposed. L-PA is associated with the pathophysiology of cerebral malaria, a severe manifestation of *Plasmodium* infection in children^53^. While previous work has demonstrated the implication of gut microbiota in pesticide detoxification in mosquitoes^54,55^, this study highlights the need to investigate how L-PA increase permethrin toxicity in *An. gambiae*.

Susceptible mosquitoes exhibited elevated mannitol levels, a metabolite known for its potential osmotic disruption properties and possibly generated by *Lactococcus lactis*^56^. but further studies are needed to understand mannitol’s impact on permethrin toxicity in *An. gambiae*.

## Methods

### Mosquito sample collections

The mosquito larvae were collected from breeding sites in Nkolondom (3°57’ 05 N and 11° 29’ 24’’E), a peri-urban area located 10 km from Yaoundé, Cameroon, where the intensive use of pesticides by farmers for their daily activity and strong pyrethroids resistance have been documented for *An. gambiae*^14^. The collection was conducted in July 2023; the larvae were brought to the Centre of Research in Infectious Diseases (CRID), where they were reared to adult F0 mosquitoes. Species ID was confirmed by PCR using the PCR molecular identification described by Bass et al.^57^.

### Insecticide susceptibility bioassay

Two to five-day-old female adult F0 mosquitoes were transferred to WHO bioassays test tubes and exposed to permethrin following an established method^58^. Four groups of 25 mosquitoes were exposed to 0.75% of permethrin-impregnated papers for 30 min and 1 hour. After the bioassays, the mosquitoes that were alive at 30 min and 1h, dead at 30 min and 1h, and unexposed controls were preserved at −80°C to avoid metabolome degradation and quench metabolic activity.

### Metabolome extraction

Metabolite extraction was performed using a method developed by Safari *et al.*^59^. Five mosquitoes from each group were individually homogenized using twenty 0.7 mm zirconia beads. Metabolites were then extracted from each sample using an ice-cold mixture of 800 µl ACN/MeOH/Acetone (8:1:1, v/v/v), the supernatant was removed, dried with nitrogen through OA-HEAT^TM^ heating system, and reconstituted in water containing 0.1% formic acid (FA).

### UHPLC-HRMS/MS conditions for untargeted metabolome

Untargeted metabolomics^60^, profiling was performed using a Q Exactive™ Hybrid Quadrupole-Orbitrap™ Mass Spectrometer coupled to an UltiMate™ 3000 UHPLC system. Chromatographic separation was achieved on an Avantor ACE Excel C18-pfp (100 × 2.1 mm, 2 µm) column. Mobile phase A was 0.1% formic acid in water, and mobile phase B was acetonitrile. Briefly, the conditions begin at 100% A for the first 3 min, followed by a gradient up to 80% B at 13 min at which it was held for 3 min before returning to starting conditions. The flow rate was 0.350 mL/min, and the column temperature was 25°C. The total run time per sample was 17 min. Heated electrospray ionization (HESI) with injection volume of 4 µl and 2 µl was used respectively in both negative and positive ion modes. HESI parameters were as follows a sheath gas flow rate of 55 arb, an auxiliary gas flow rate of 10 arb, a sweeping gas flow rate of 1 arb, and a capillary temperature of 325°C. The spray voltage was 3.00 kV for negative mode and 3.50 kV for positive mode. The S-lens RF level was set to 30. Full scan data (m/z 70-1000) were acquired at a resolving power of 35,000. For tandem mass spectrometry (MS/MS) with biological samples, an isolation window of 2.0 m/z was used. Automatic gain control (AGC) target was 5×10⁵, with a maximum injection time (IT) of 75 ms. The normalized collision energy (NCE) was 40, and data was acquired at a resolving power of 17,500. For analysis of chemical standards, the same parameters were used.

### Data processing and Statistical analysis

Raw UHPLC-HRMS/MS data were converted to mzXML using Raw Converter and subsequently processed with MZmine 2.53^61,62^. Statistical analysis was performed using MetaboAnalyst 6.0.^63^ and GraphPad Prism 10. Data filtering with variance filter by standard deviation and abundance filter by mean intensity value with RDS greater than 25%. Normalization was grouped into 3 categories, normalization by sum, data log transformation (base 10) and data auto scaling, ANOVA, post-hoc Fisher’s tests with false discovery rate (FDR)-corrected p-value threshold=0.05.

## Conclusion

Through comprehensive untargeted metabolomic profiling, we have characterized the metabolomic profile of permethrin resistance of field-derived *An. gambiae* from Cameroon. We observed metabolite signals that align with P450 monooxygenase-mediated detoxification, target site resistance mechanisms, and identified metabolic shifts indicative of enhanced energy production in resistant mosquitoes. We further revealed that NAA may influence neuronal energy homeostasis, which may modulate toxicity of permethrin. Both XA and 3-hydroxykynurenine, derived from tryptophan catabolism, as well as L-pipecolic acid, a catabolite of lysine, are associated with permethrin toxicity. These findings improve our understanding of the role of metabolism in *An*. *gambiae* permethrin-resistance and compels further interrogation of the role of these pathways in permethrin cross-resistance.

## Acknowledgments

This study was supported, in part, by U19AI181594 from the US National Institutes of Health to R.R.D, C.S.W, and T.J.G., and a Bill and Melinda Gates foundation project INV-006003 grant to C.S.W.

## Author contribution

C.S.W. and T. J. G. conceived and designed the study; S.V.D., M.K., and C.S.W. performed the field collection and resistance bioassays. S.V.D, M.K., SSI., and T. J. G. performed the sample preparation for UHPLC/HRMS/MS. UHPLC/HRMS/MS data collection and data processing were performed by T. J. G., M.W.C. and S.V.D. MS/MS data analysis and confirmation was done by T. J. G. with the assistance of M.W.C. and S.V.D. The paper was written by S.V.D. with assistance from S.W.T., T. J. G., R.R.D., S.S.I. and M.W.C. T.J.G., R.R.D., and C.S.W. edited the last version. All the authors read and approved the final draft of the manuscript.

## Competing Information

The authors declare no competing interest.

## Additional information

Supplementary information is available for this manuscript.

