## Supplemental data for "Metabolomic changes underpinning permethrin resistance in the *Anopheles gambiae* malaria vector from Cameroon"

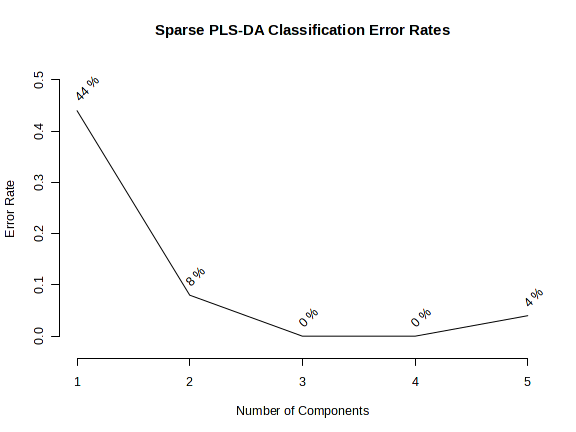

**Figure S1.** Performance of Sparse partial least squares-discriminant analysis (sPLS-DA)

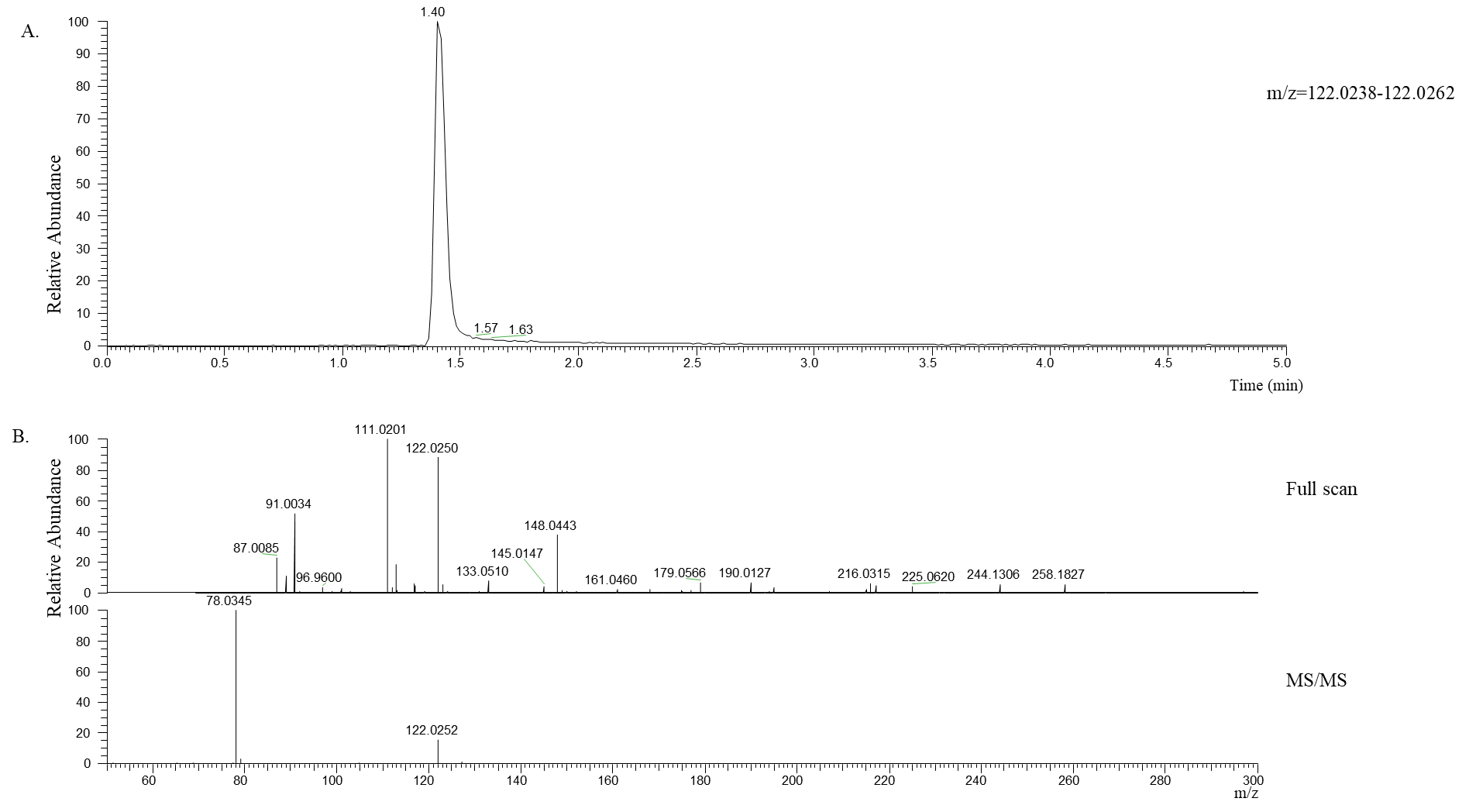

**Figure S2.**  Negative ionization for the identification of nicotinate.  Top, extracted ion chromatogram for m/z 122.0250 (5 ppm). Middle, full scan mass spectrum from retention time 1.40 min.  Bottom, tandem mass spectrum (MS/MS) for m/z 122.0250 (1.2 amu isolation window).

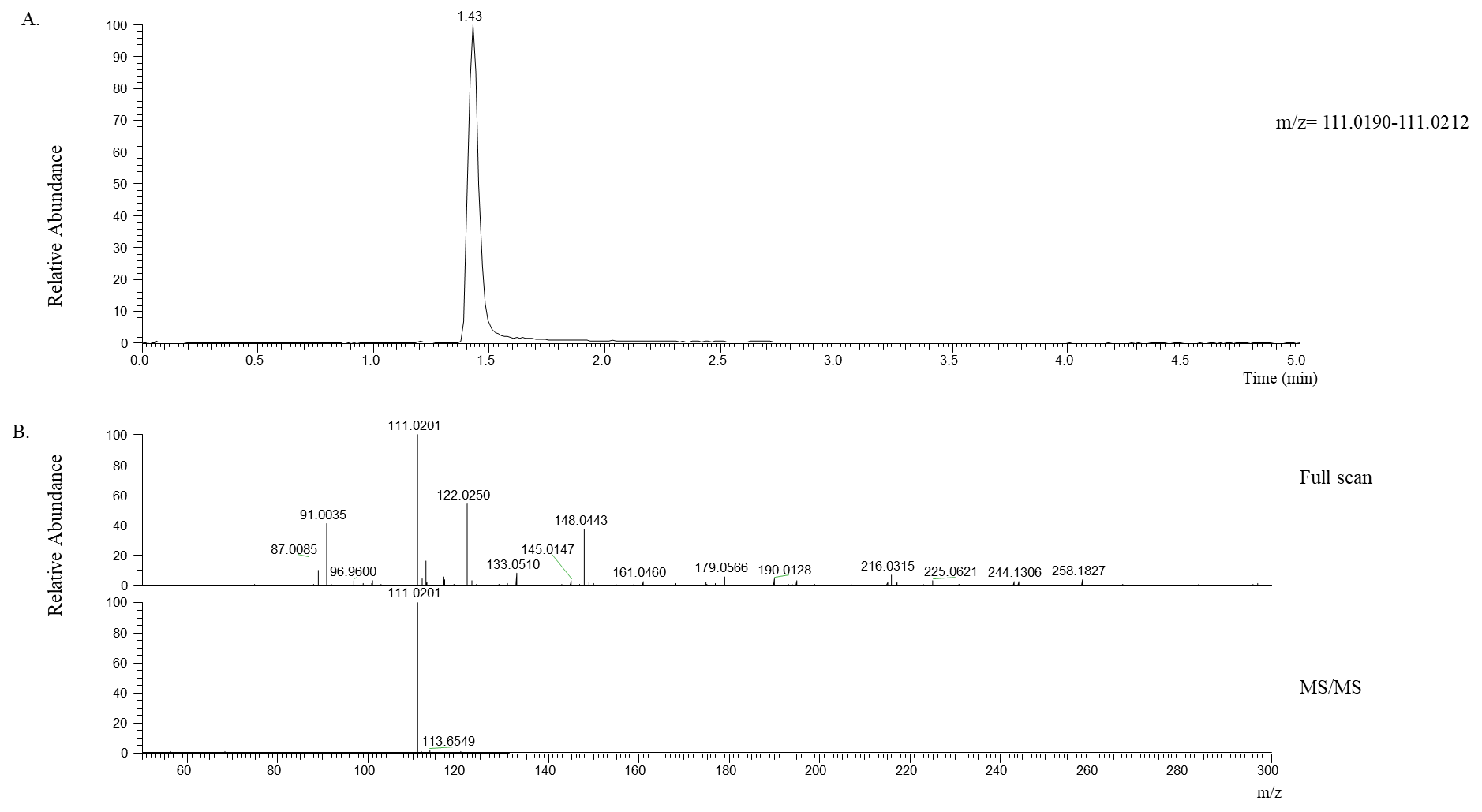

**Figure S3.**  Negative ionization for the identification of uracil.  Top, extracted ion chromatogram for m/z 111.0201 (5 ppm). Middle, full scan mass spectrum from retention time 1.43 min.  Bottom, tandem mass spectrum (MS/MS) for m/z 111.0201 (1.2 amu isolation window).

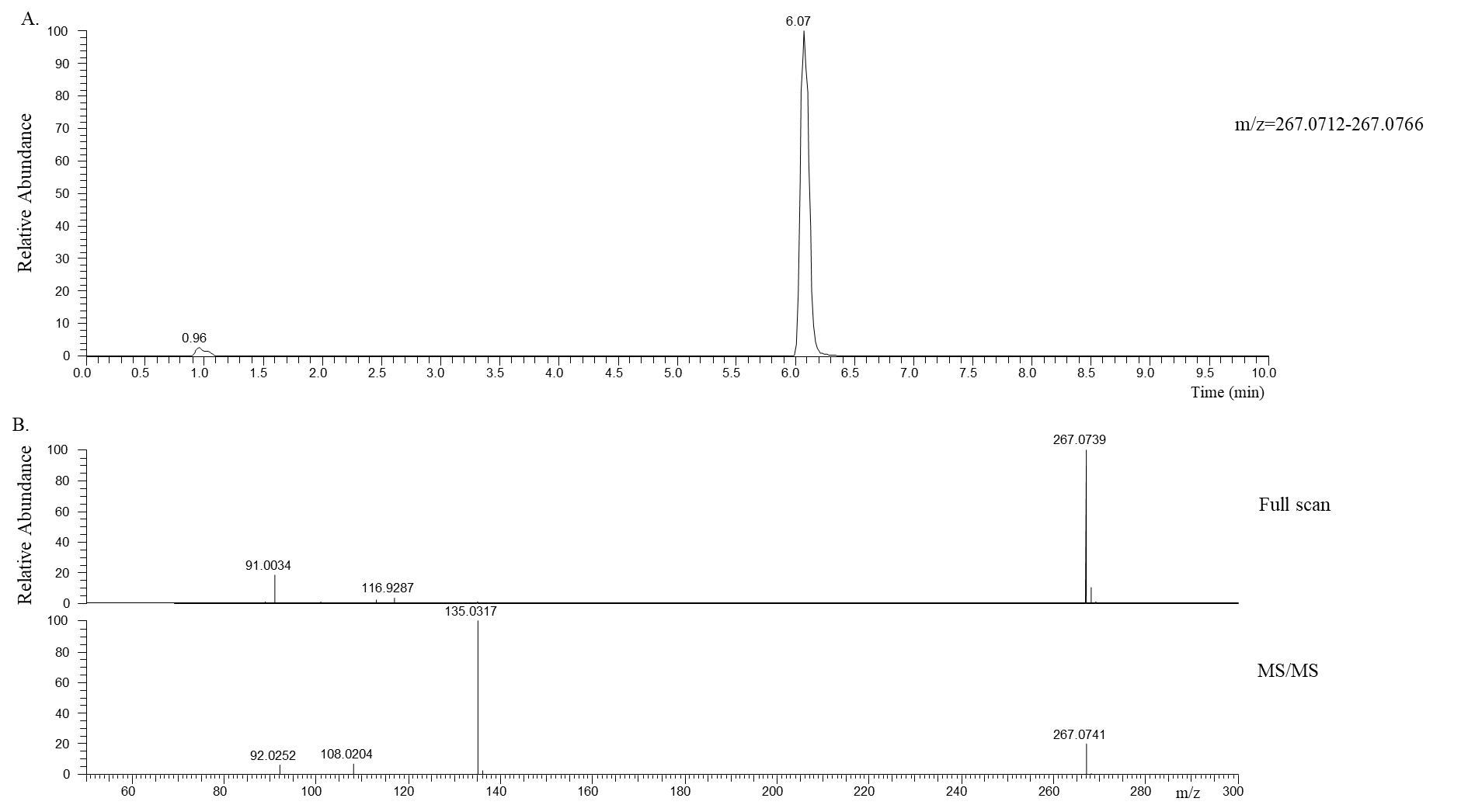

**Figure S4.**  Negative ionization for the identification of inosine.  Top, extracted ion chromatogram for m/z 267.0739 (5 ppm). Middle, full scan mass spectrum from retention time 6.07 min.  Bottom, tandem mass spectrum (MS/MS) for m/z 267.0741 (1.2 amu isolation window).

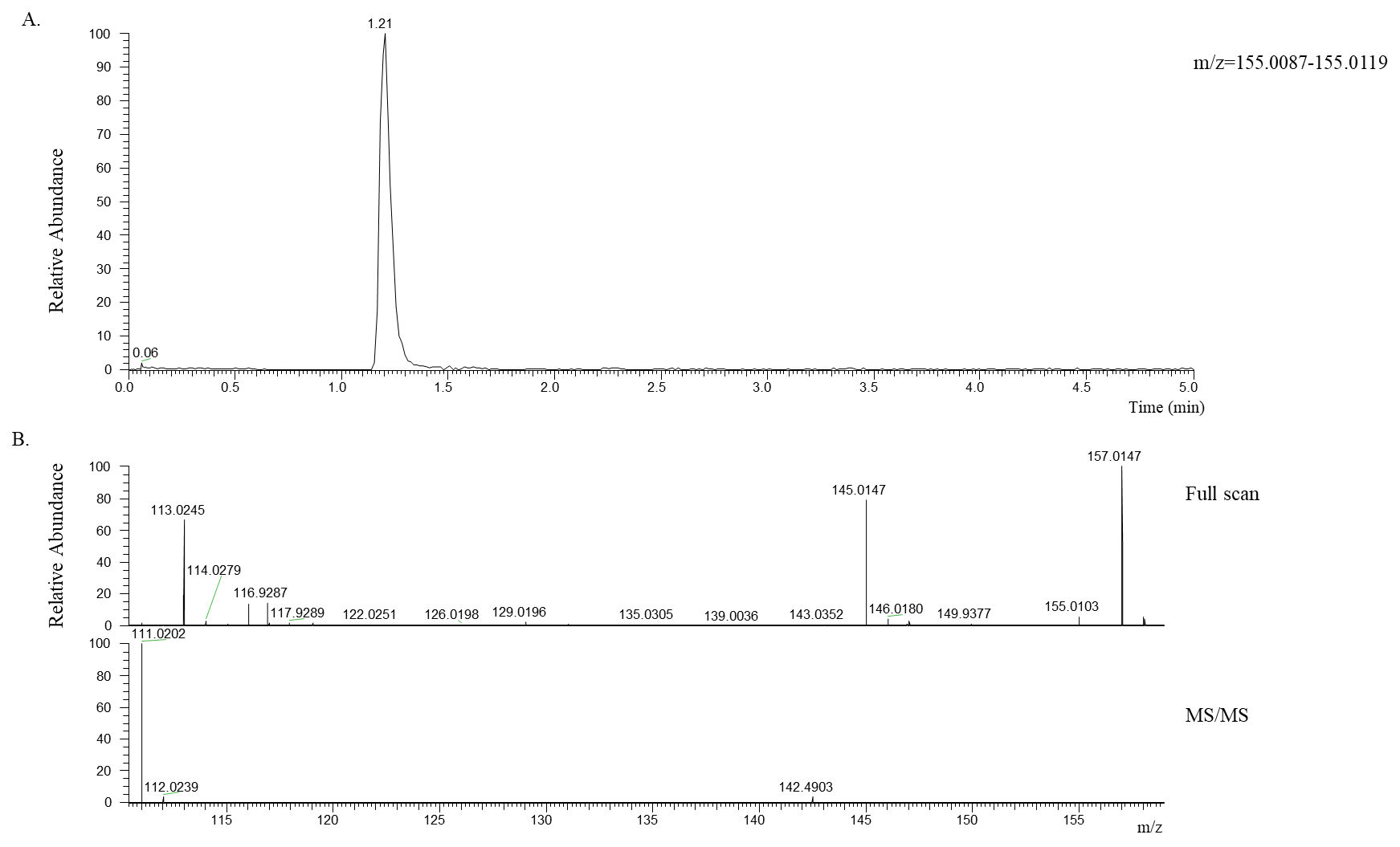

**Figure S5.**  Positive ionization for the identification of orotate.  Top, extracted ion chromatogram for m/z 155.0087 (5 ppm). Middle, full scan mass spectrum from retention time 1.21 min.  Bottom, tandem mass spectrum (MS/MS) for m/z 155.0103 (1.2 amu isolation window).

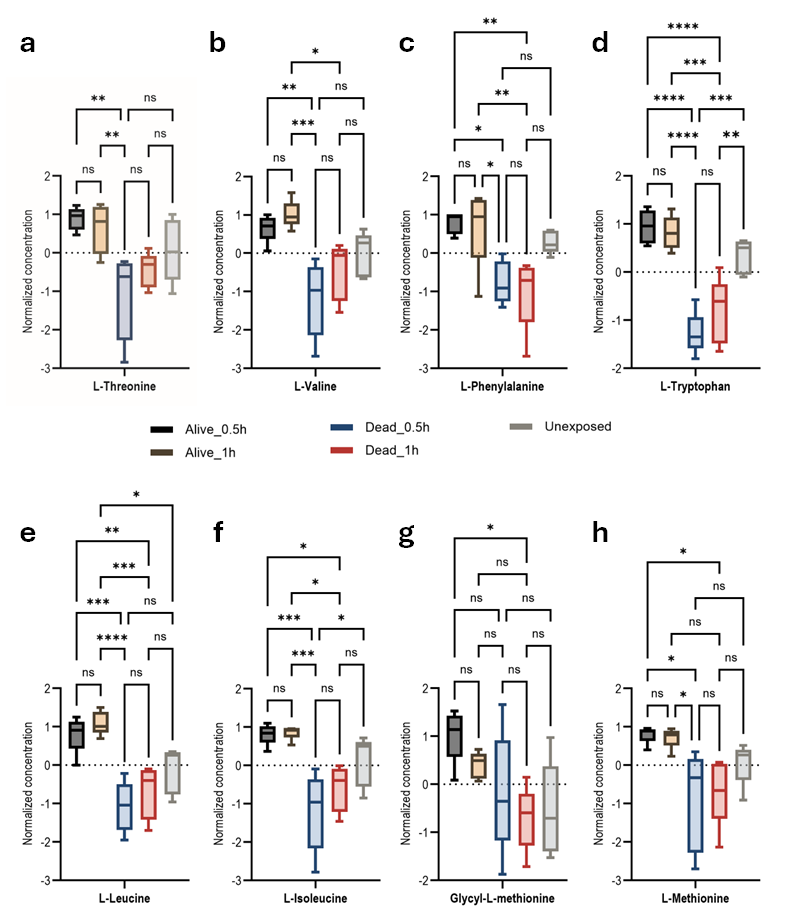

**Figure S6**. Summary of amino acid profile-based *An. gambiae* permethrin phenotypes. Anova, post-hoc Fisher with p-value p-value threshold=0.05, statistical annotation: ns, non-significant; threonine, **p-value=0.002; L-Valine, *p-value=0.014, **p-value=0.00199, ***p-value=0.0003; L-phenylalanine, *p-value=0.0174, **p-value=0.0052; L-tryptophan, **p-value=0.0059, ***p-value=0.0001, ****p-value˂0.0001; L-leucine, *p-value=0.0216, **p-value=0.0034, ***p-value=0.0003, ****p-value˂0.0001; L-isoleucine, *p-value=0.0151, ***p-value=0.0005; L-methionine, *p-value=0.0155; Glycyl-methionine *p-value=0.012.

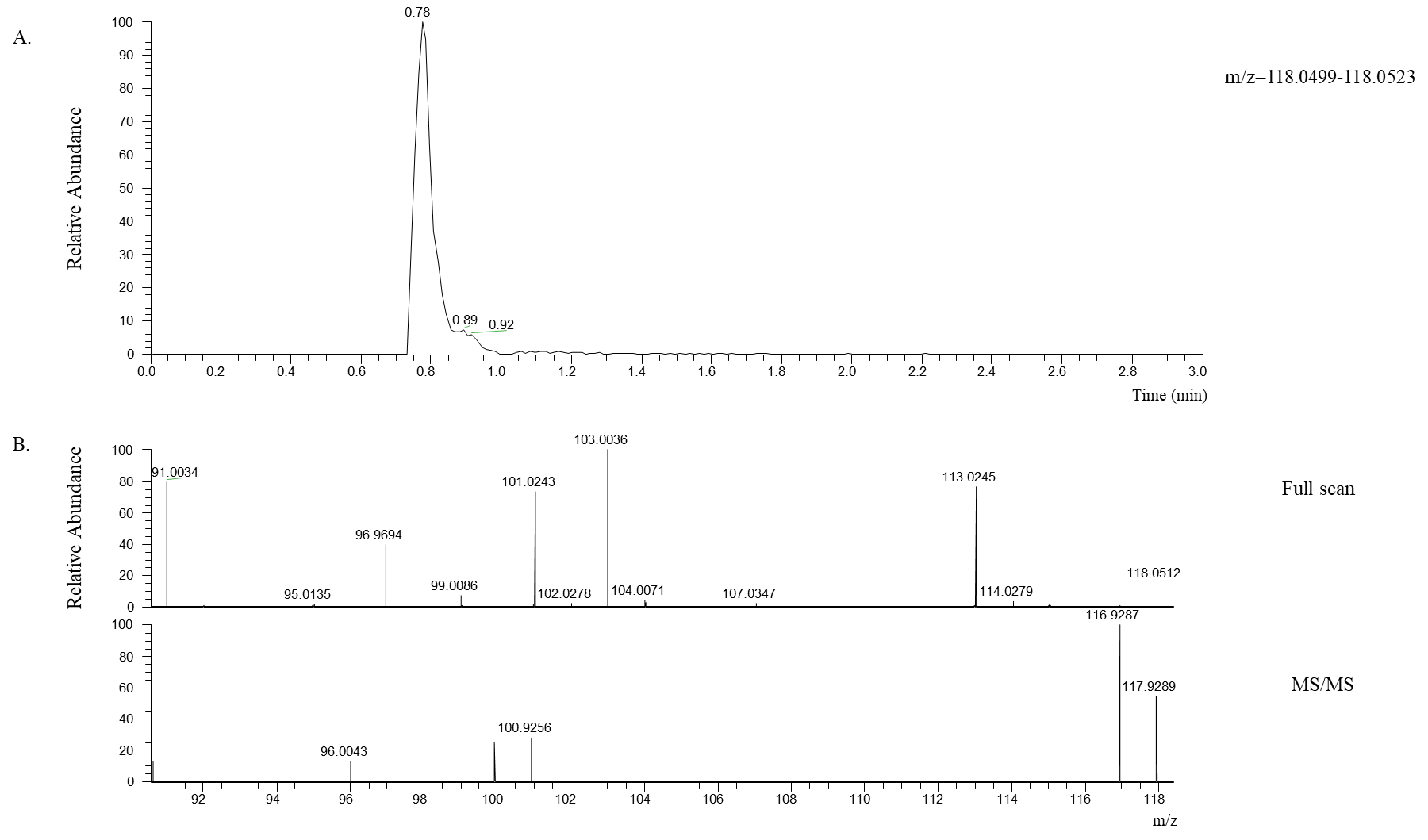

**Figure S7.**  Negative ionization for the identification of L-threonine.  Top, extracted ion chromatogram for m/z 118.0512 (5 ppm). Middle, full scan mass spectrum from retention time 0.78 min.  Bottom, tandem mass spectrum (MS/MS) for m/z 118.0512 (1.2 amu isolation window).

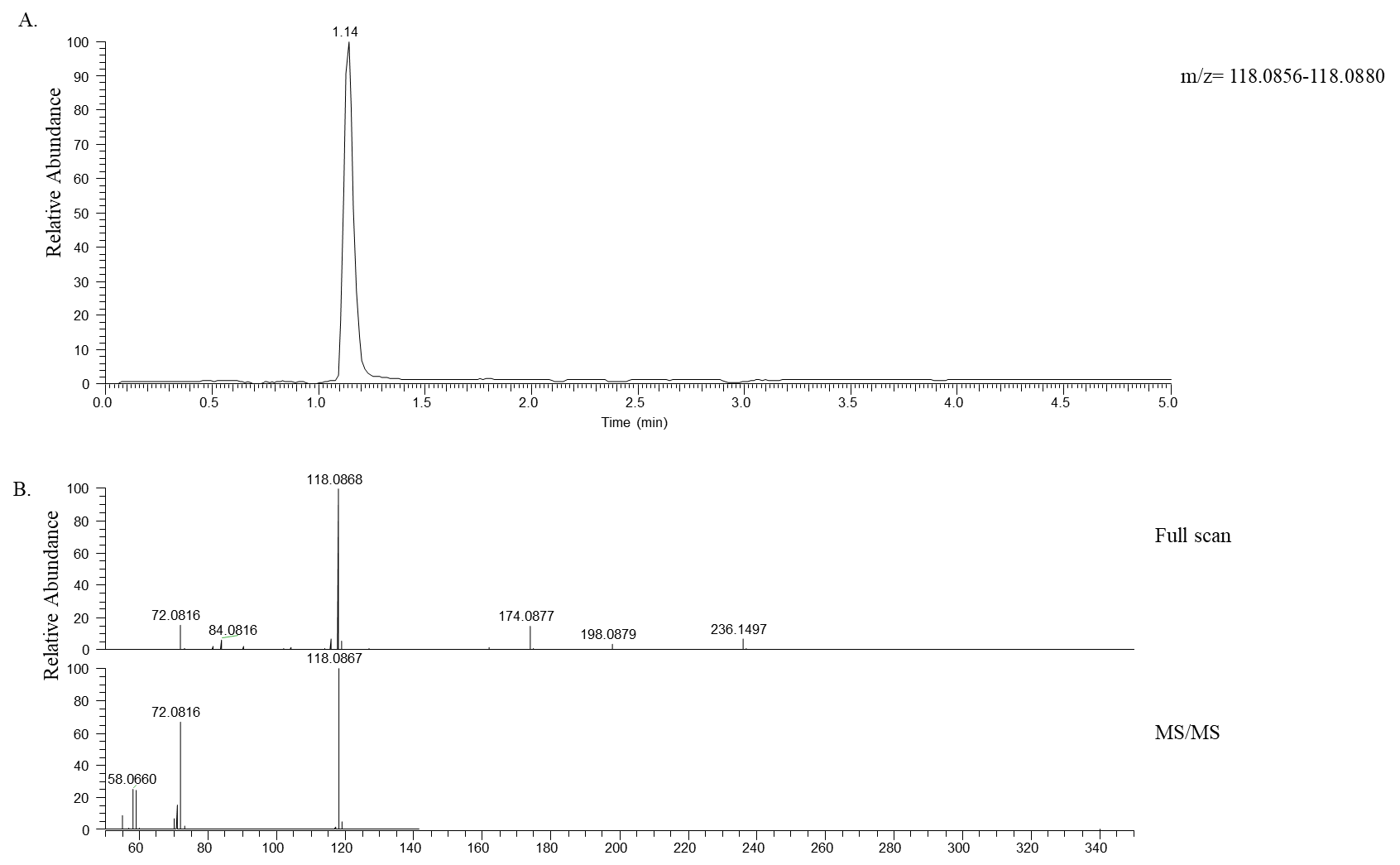

**Figure S8.**  Positive ionization for the identification of L-valine.  Top, extracted ion chromatogram for m/z 118.0868 (5 ppm). Middle, full scan mass spectrum from retention time 1.14 min.  Bottom, tandem mass spectrum (MS/MS) for m/z 118.0867 (1.2 amu isolation window).

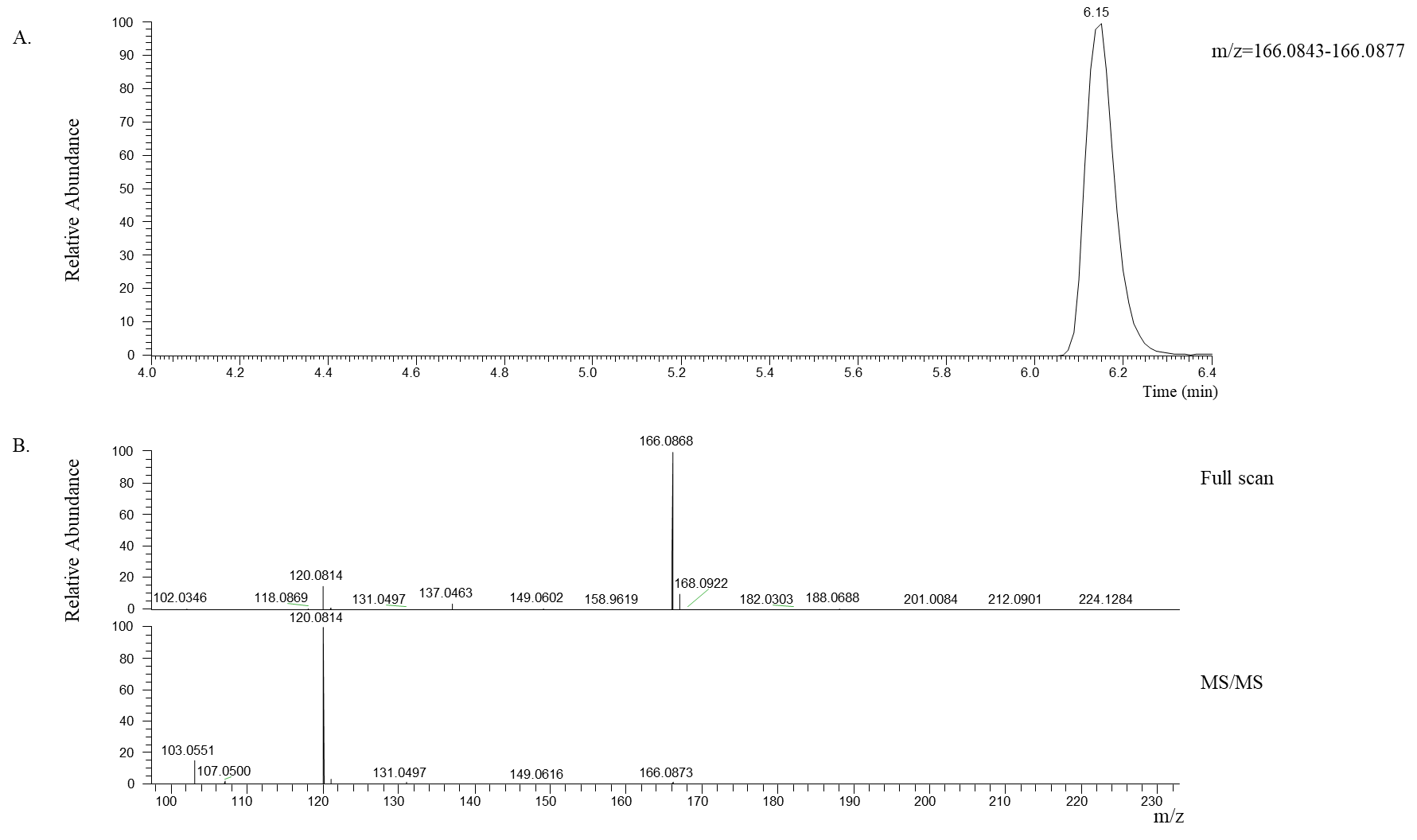

**Figure S9.**  Positive ionization for the identification of L-phenylalanine.  Top, extracted ion chromatogram for m/z 166.0868 (5 ppm). Middle, full scan mass spectrum from retention time 6.15 min.  Bottom, tandem mass spectrum (MS/MS) for m/z 166.0868 (1.2 amu isolation window).

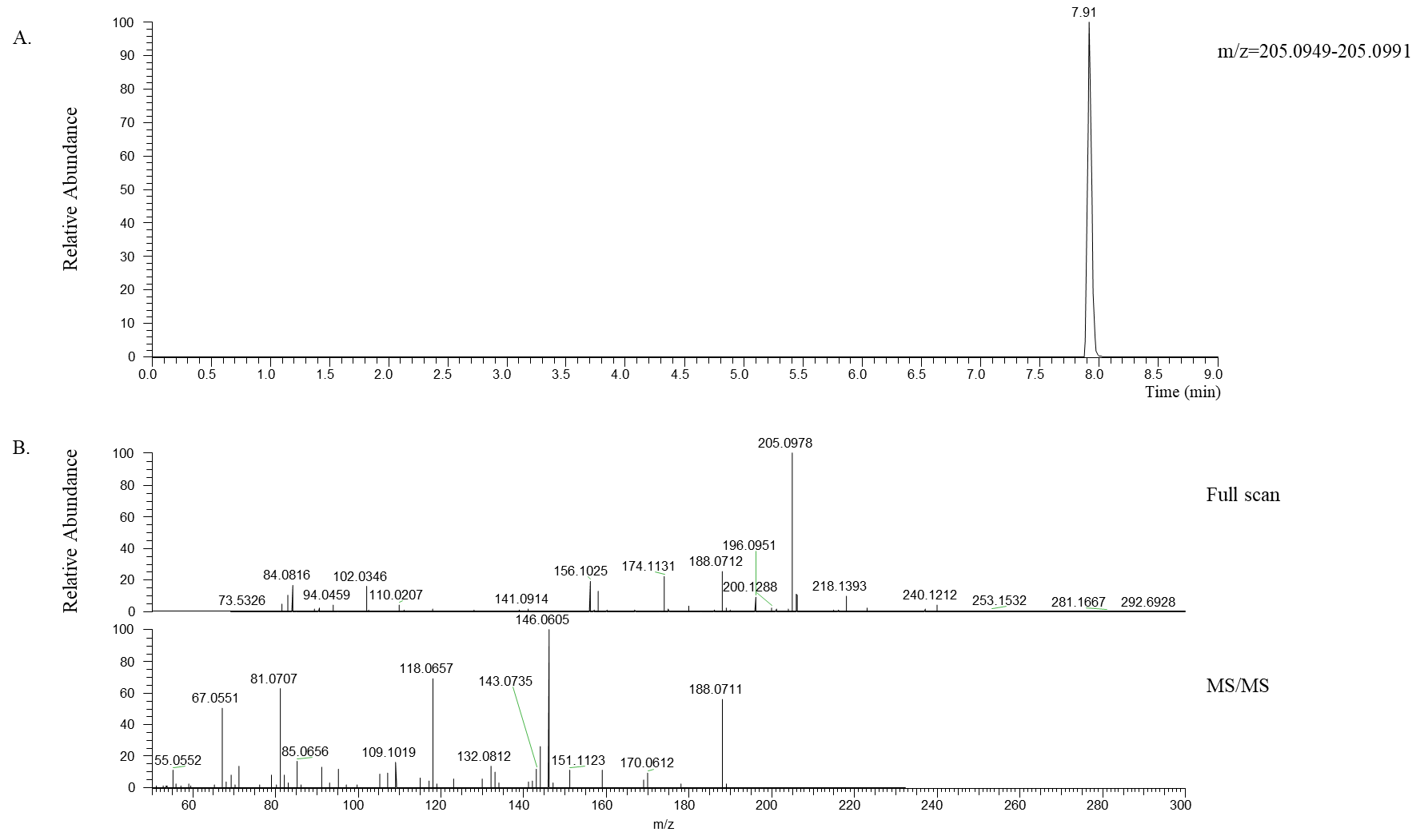

**Figure S10.**  Negative ionization for the identification of L-tryptophane.  Top, extracted ion chromatogram for m/z 205.0978 (5 ppm). Middle, full scan mass spectrum from retention time 7.91 min.  Bottom, tandem mass spectrum (MS/MS) for m/z 205.0978 (1.2 amu isolation window).

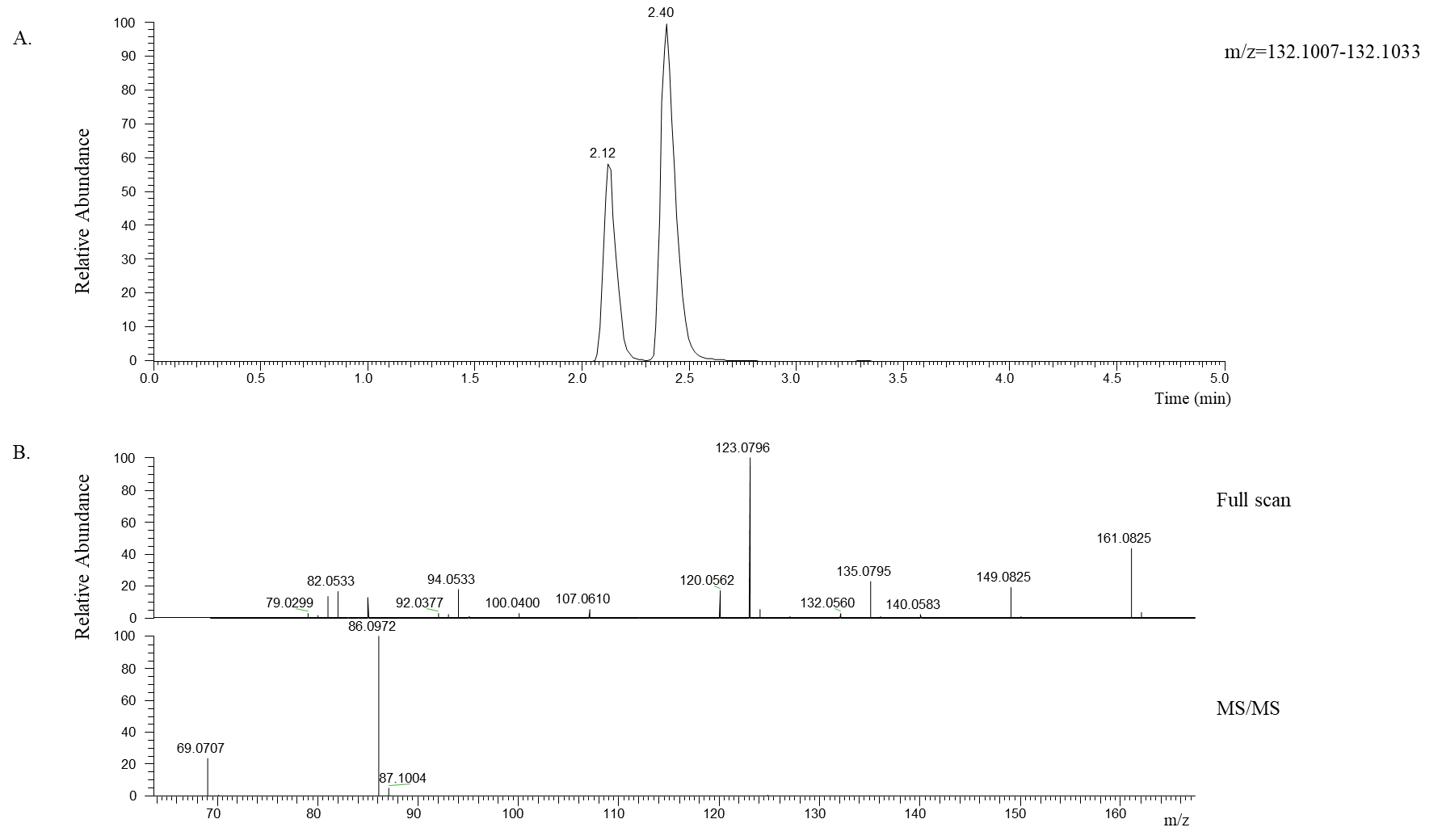

**Figure S11.**  Positive ionization for the identification of L-leucine.  Top, extracted ion chromatogram for m/z 132.0560 (5 ppm). Middle, full scan mass spectrum from retention time 2.40 min.  Bottom, tandem mass spectrum (MS/MS) for m/z 205.0978 (1.2 amu isolation window).

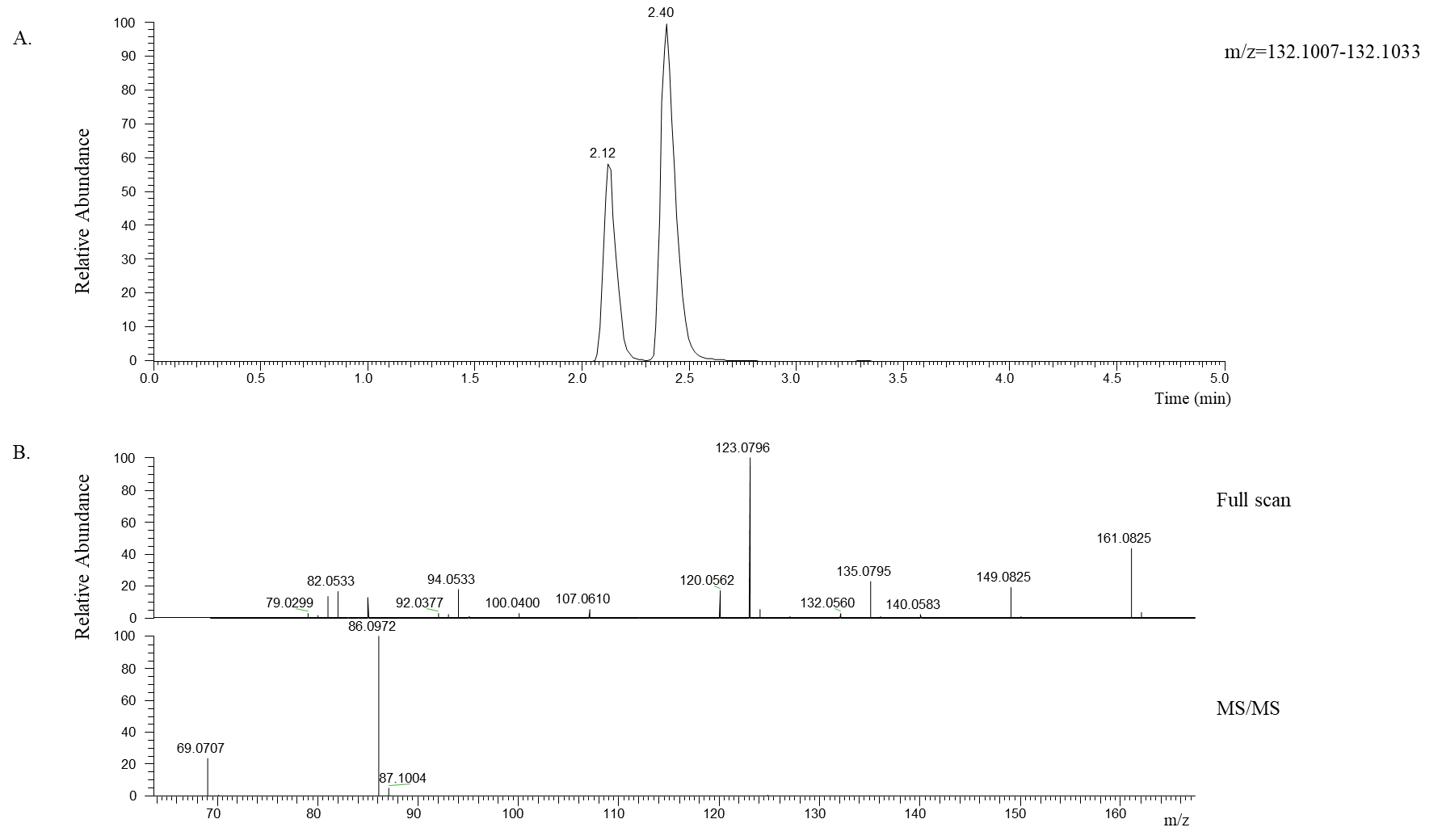

**Figure S12.**  Positive ionization for the identification of L-leucine.  Top, extracted ion chromatogram for m/z 132.0560 (5 ppm). Middle, full scan mass spectrum from retention time 2.12 min.  Bottom, tandem mass spectrum (MS/MS) for m/z 205.0978 (1.2 amu isolation window).

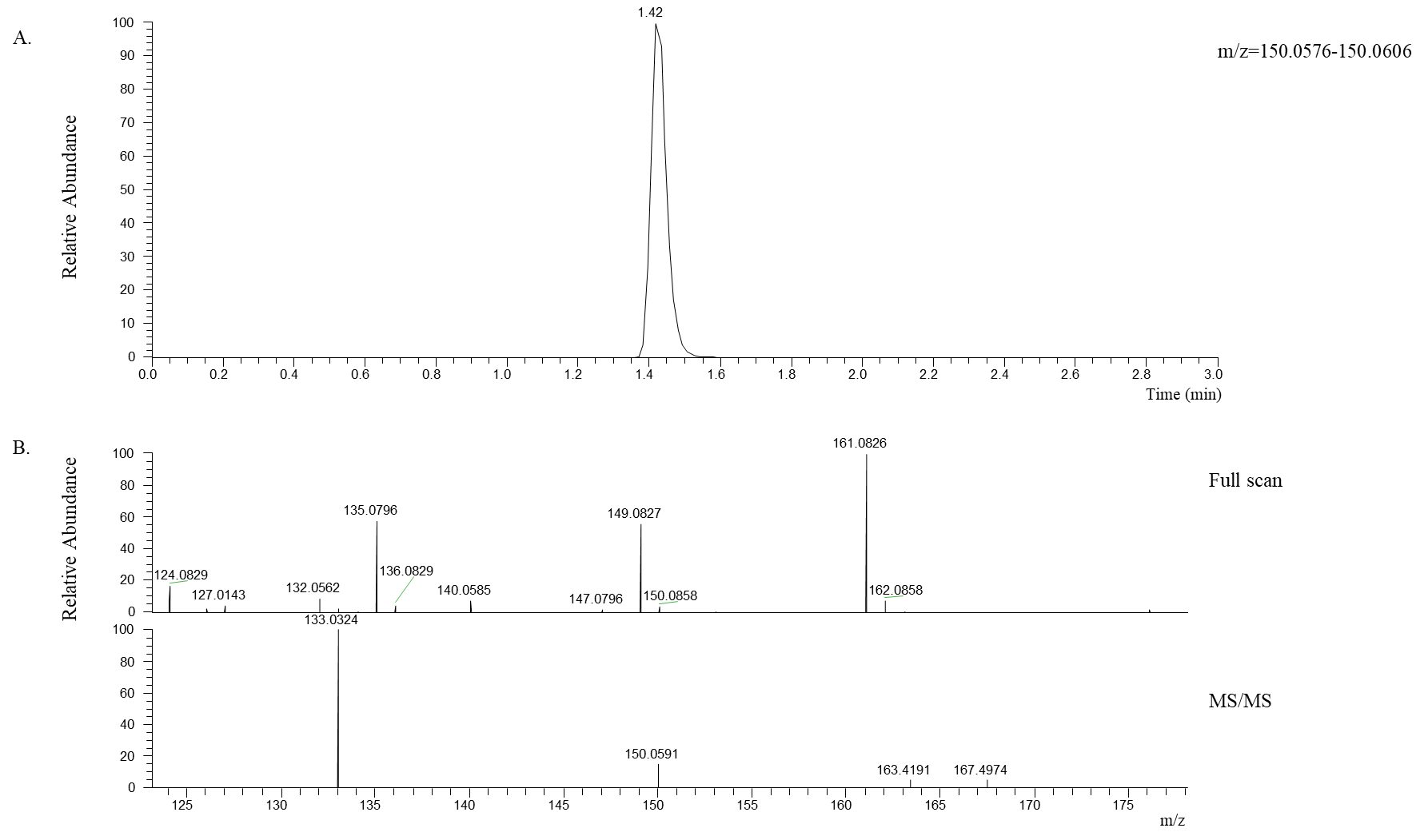

**Figure S13.**  Positive ionization for the identification of L-methionine.  Top, extracted ion chromatogram for m/z 150.0858 (5 ppm). Middle, full scan mass spectrum from retention time 1.42 min.  Bottom, tandem mass spectrum (MS/MS) for m/z 150.0591 (1.2 amu isolation window).

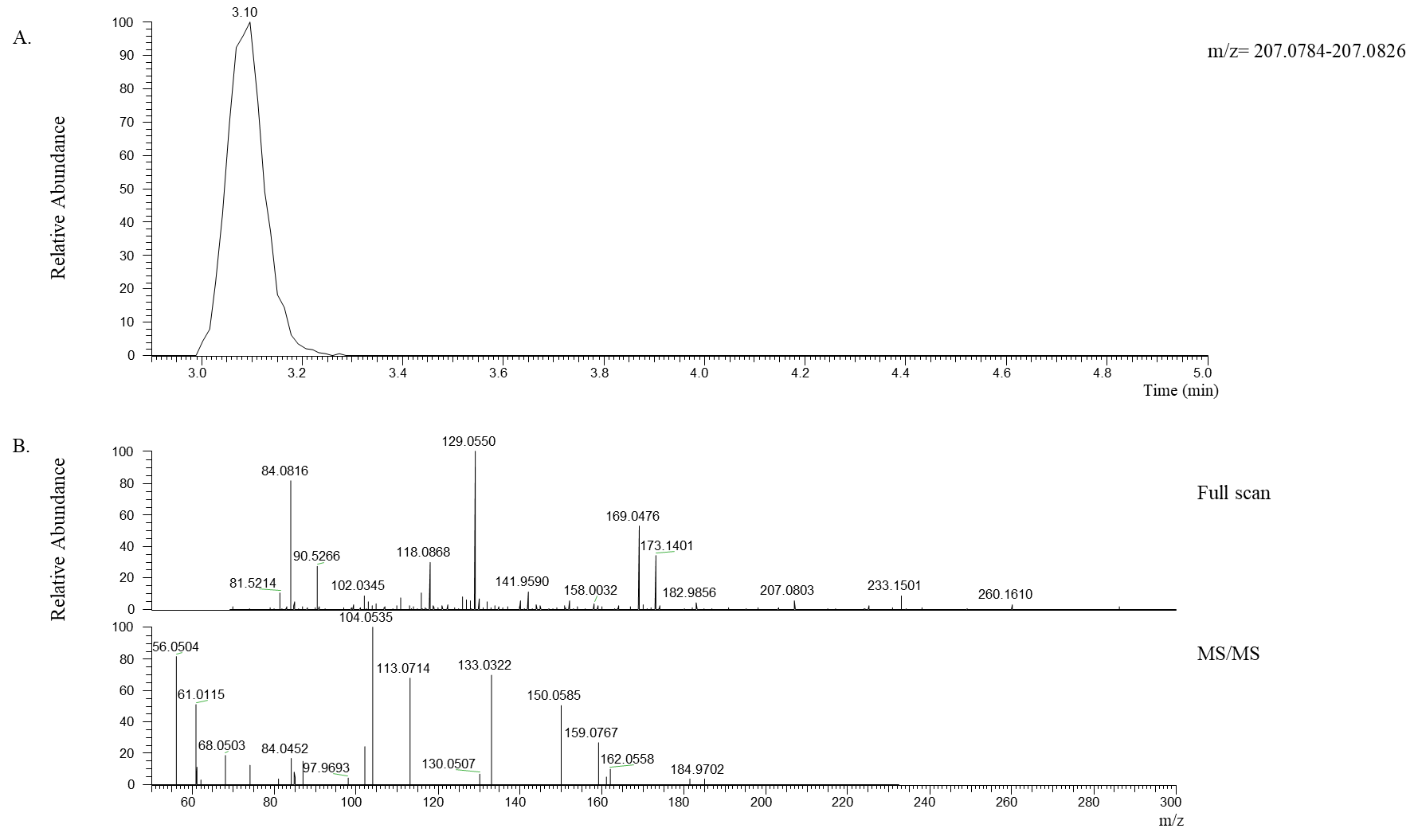

**Figure S14.**  Positive ionization for the identification of glycyl-L-methionine.  Top, extracted ion chromatogram for m/z 207.0803 (5 ppm). Middle, full scan mass spectrum from retention time 3.10 min.  Bottom, tandem mass spectrum (MS/MS) for m/z 207.0803 (1.2 amu isolation window).

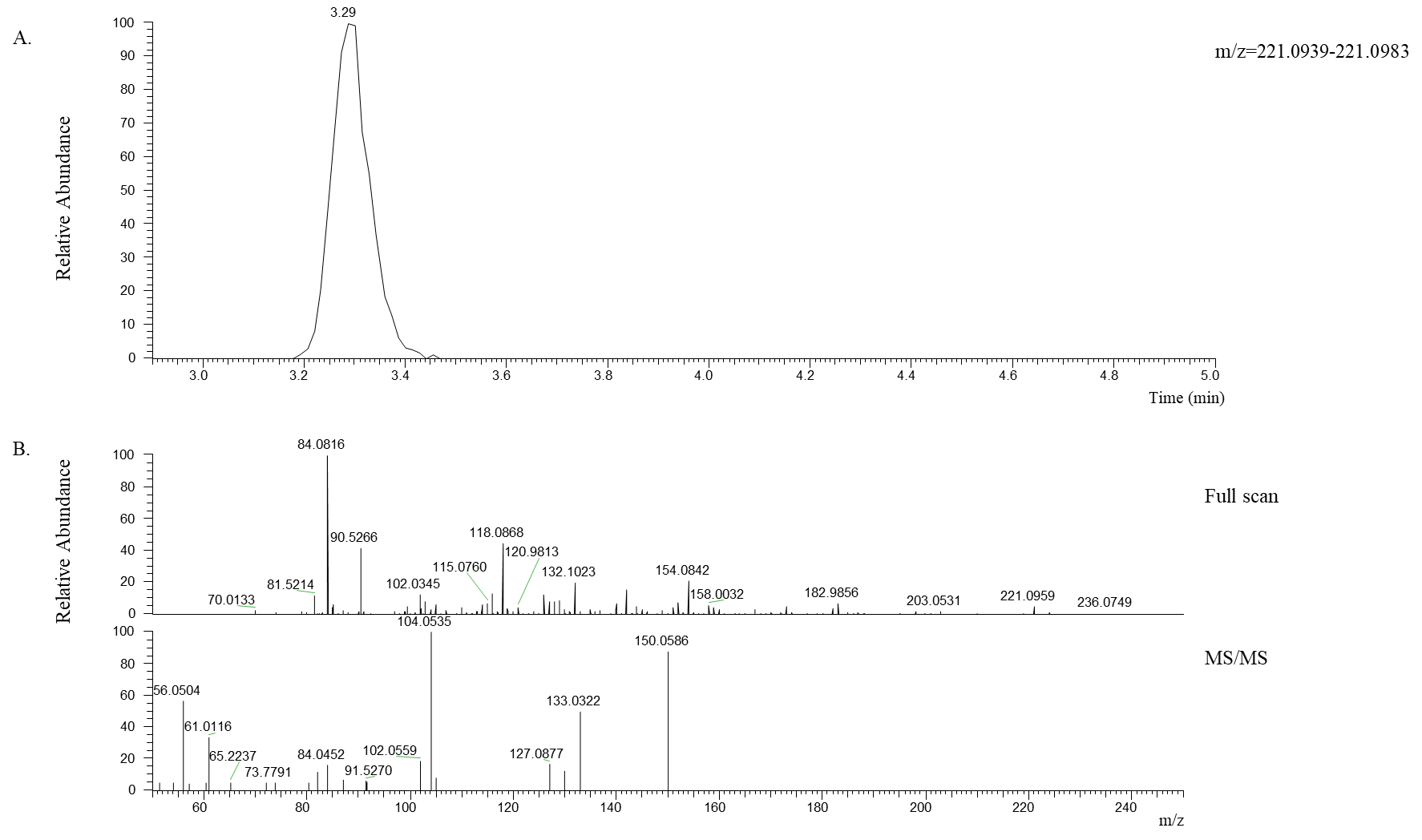

**Figure S15.**  Positive ionization for the identification of alanyl-L-methionine.  Top, extracted ion chromatogram for m/z 221.0959 (5 ppm). Middle, full scan mass spectrum from retention time 3.29 min.  Bottom, tandem mass spectrum (MS/MS) for m/z 221.0959 (1.2 amu isolation window).

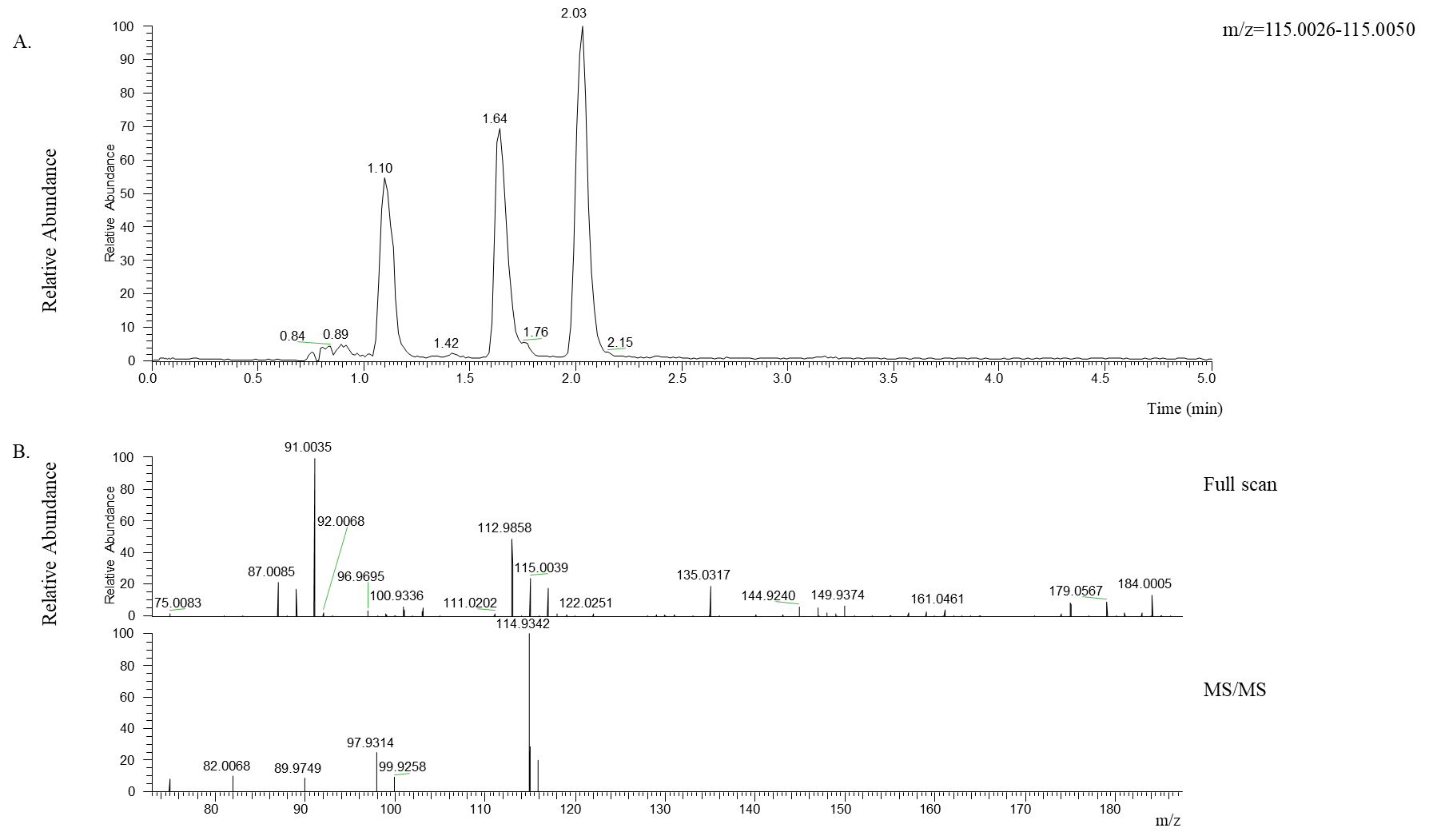

**Figure S16.**  Negative ionization for the identification of fumarate.  Top, extracted ion chromatogram for m/z 115.0039 (5 ppm). Middle, full scan mass spectrum from retention time 2.03 min.  Bottom, tandem mass spectrum (MS/MS) for m/z 115.0039 (1.2 amu isolation window).

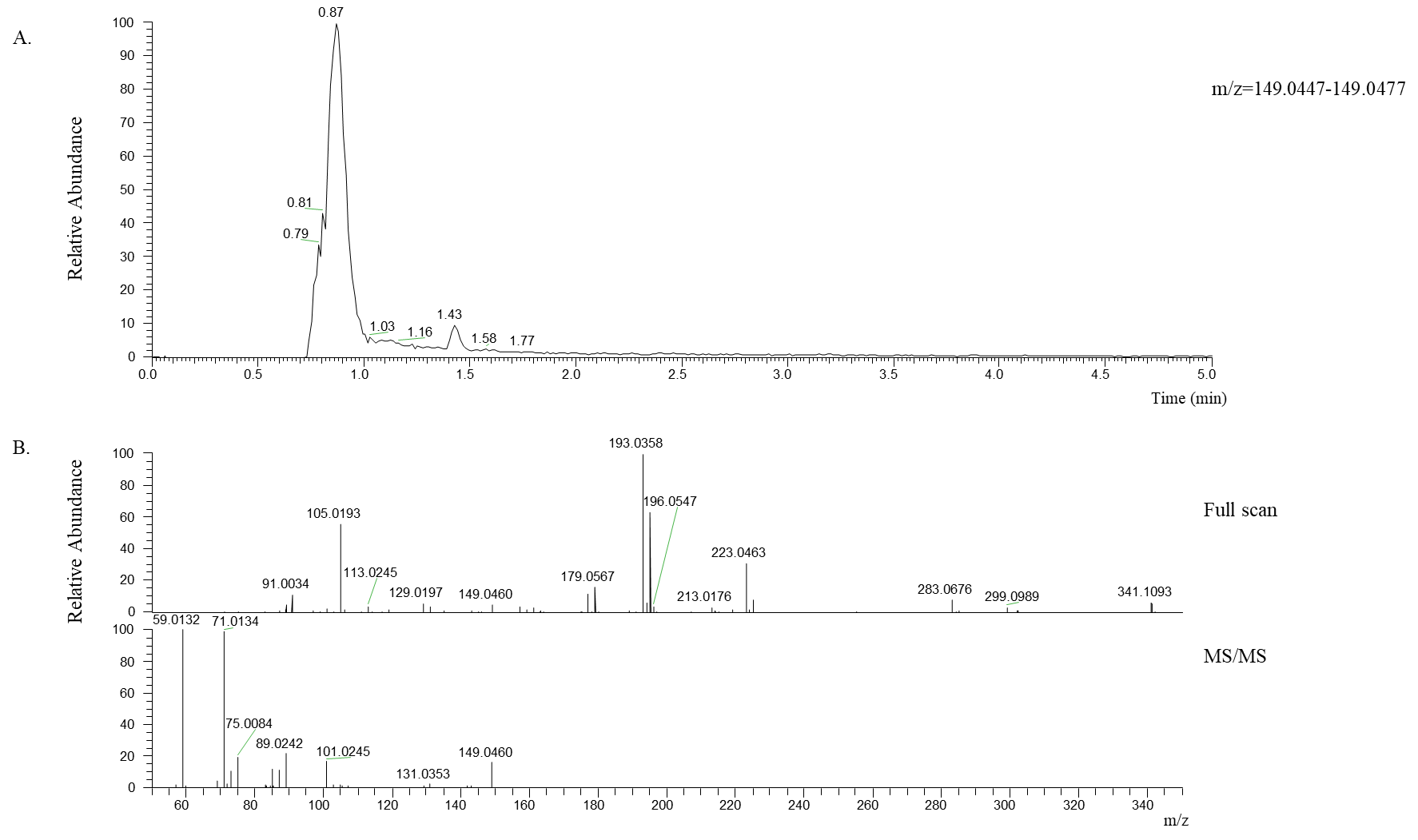

**Figure S17.**  Negative ionization for the identification of fumarate.  Top, extracted ion chromatogram for m/z 149.0460 (5 ppm). Middle, full scan mass spectrum from retention time 0.87 min.  Bottom, tandem mass spectrum (MS/MS) for m/z 149.0460 (1.2 amu isolation window).

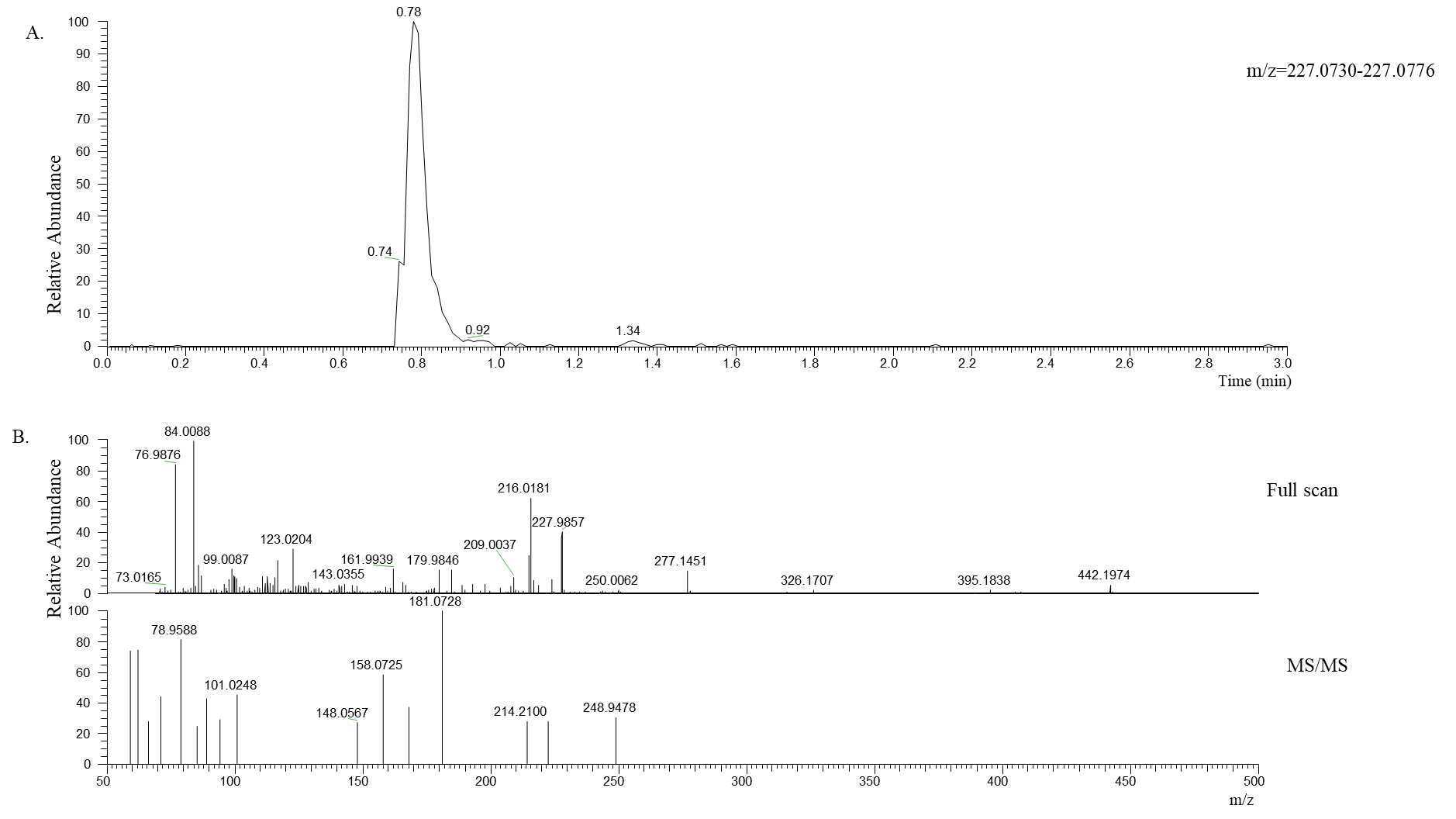

**Figure S18.**  Negative ionization for the identification of mannitol.  Top, extracted ion chromatogram for m/z 227.9857 (5 ppm). Middle, full scan mass spectrum from retention time 0.78 min.  Bottom, tandem mass spectrum (MS/MS) for m/z 227.9857 (1.2 amu isolation window).

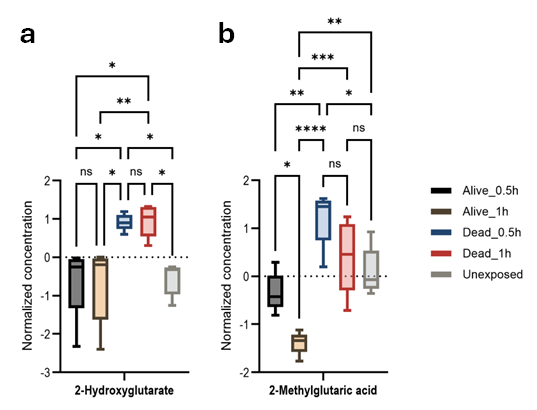

**Figure S19**: Hypoxia markers associated with permethrin neurotoxic effects in *An. gambiae* permethrin phenotypes. Anova, post-hoc Fisher with p-value p-value threshold=0.05, statistical.

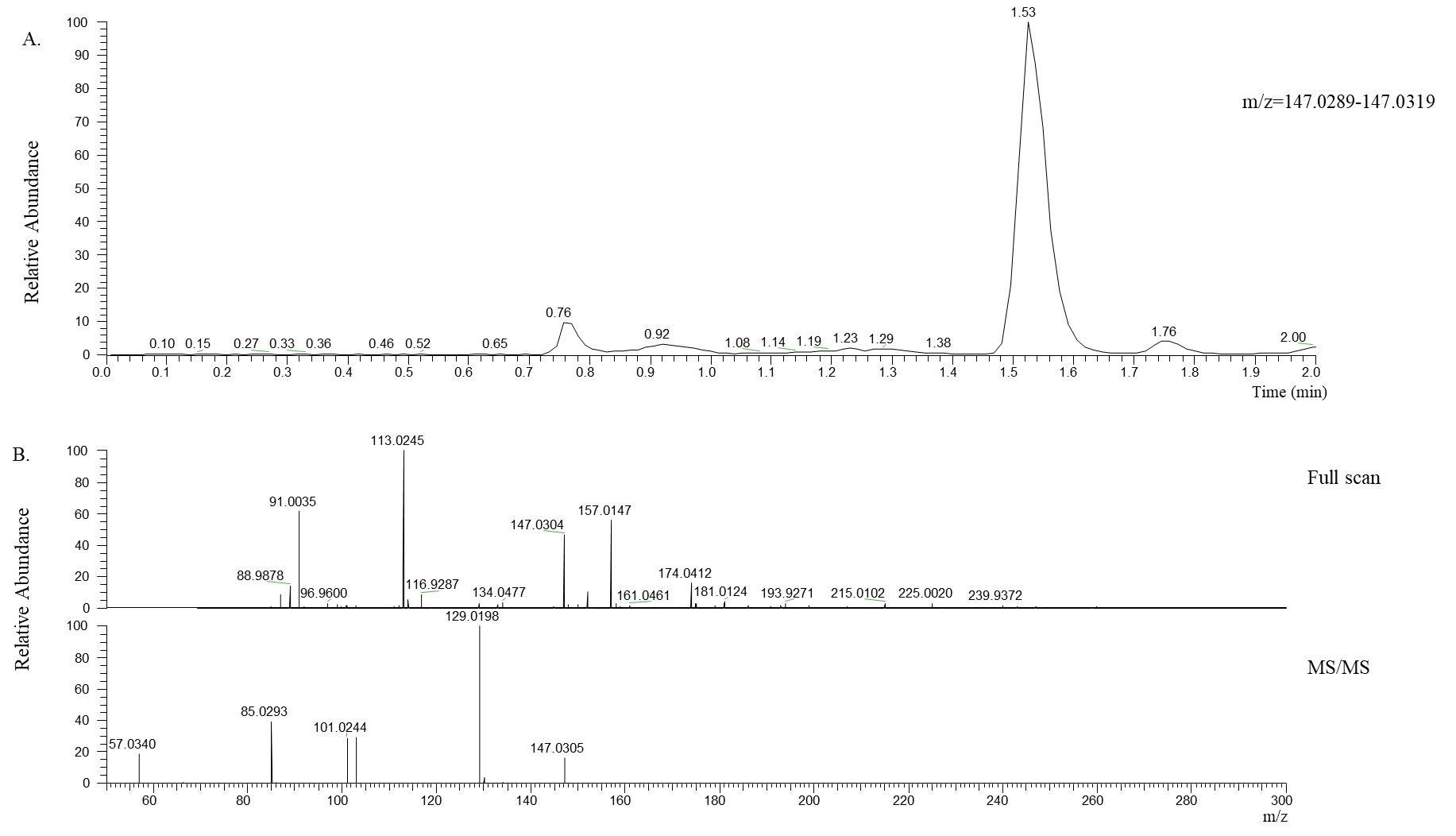

**Figure S20.**  Negative ionization for the identification of 2-hydroxyglutarate.  Top, extracted ion chromatogram for m/z 147.0304 (5 ppm). Middle, full scan mass spectrum from retention time 1.53 min.  Bottom, tandem mass spectrum (MS/MS) for m/z 147.0304 (1.2 amu isolation window).

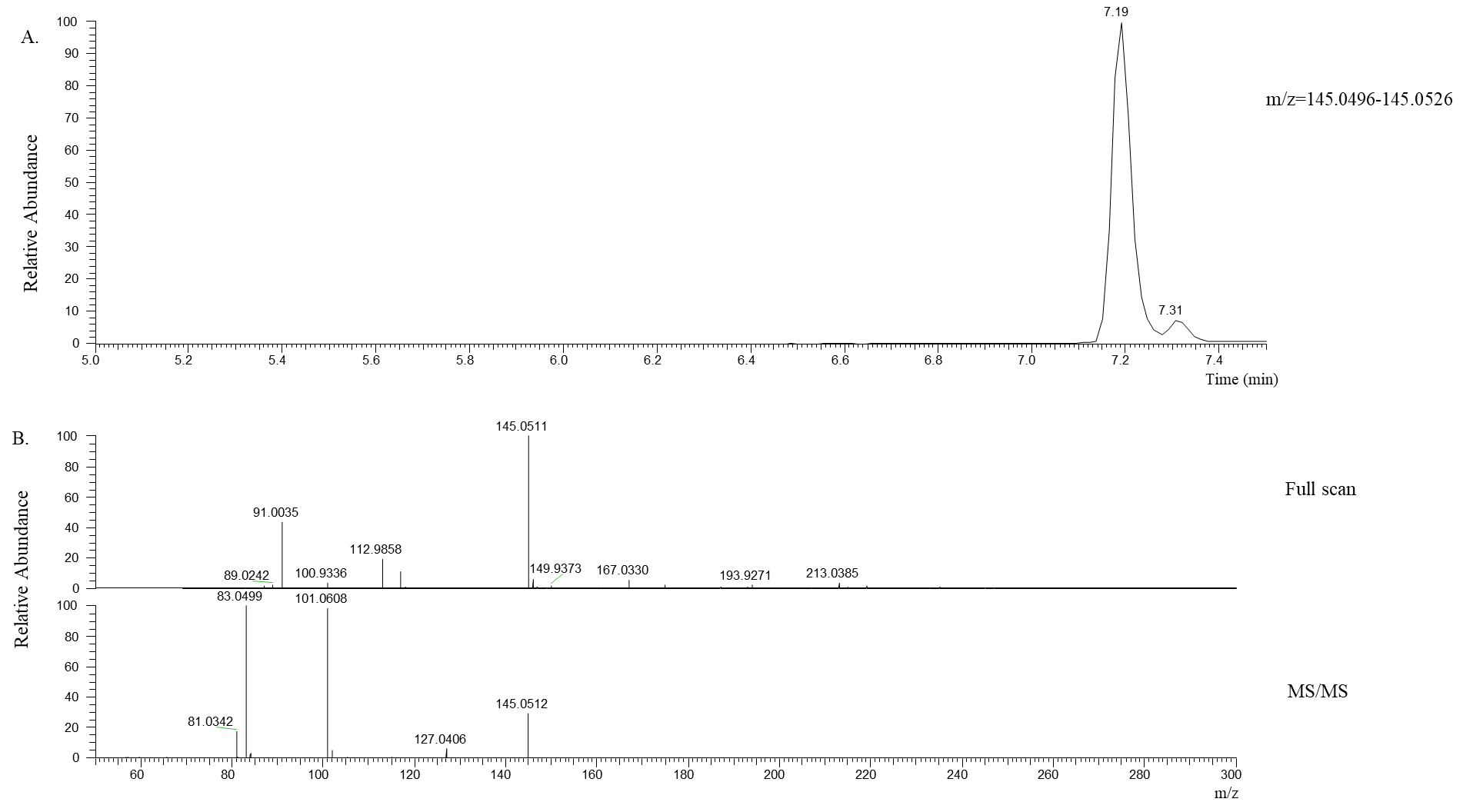

**Figure S21.**  Negative ionization for the identification of 2-methylglutarate.  Top, extracted ion chromatogram for m/z 145.0511 (5 ppm). Middle, full scan mass spectrum from retention time 7.19 min.  Bottom, tandem mass spectrum (MS/MS) for m/z 145.0511 (1.2 amu isolation window).

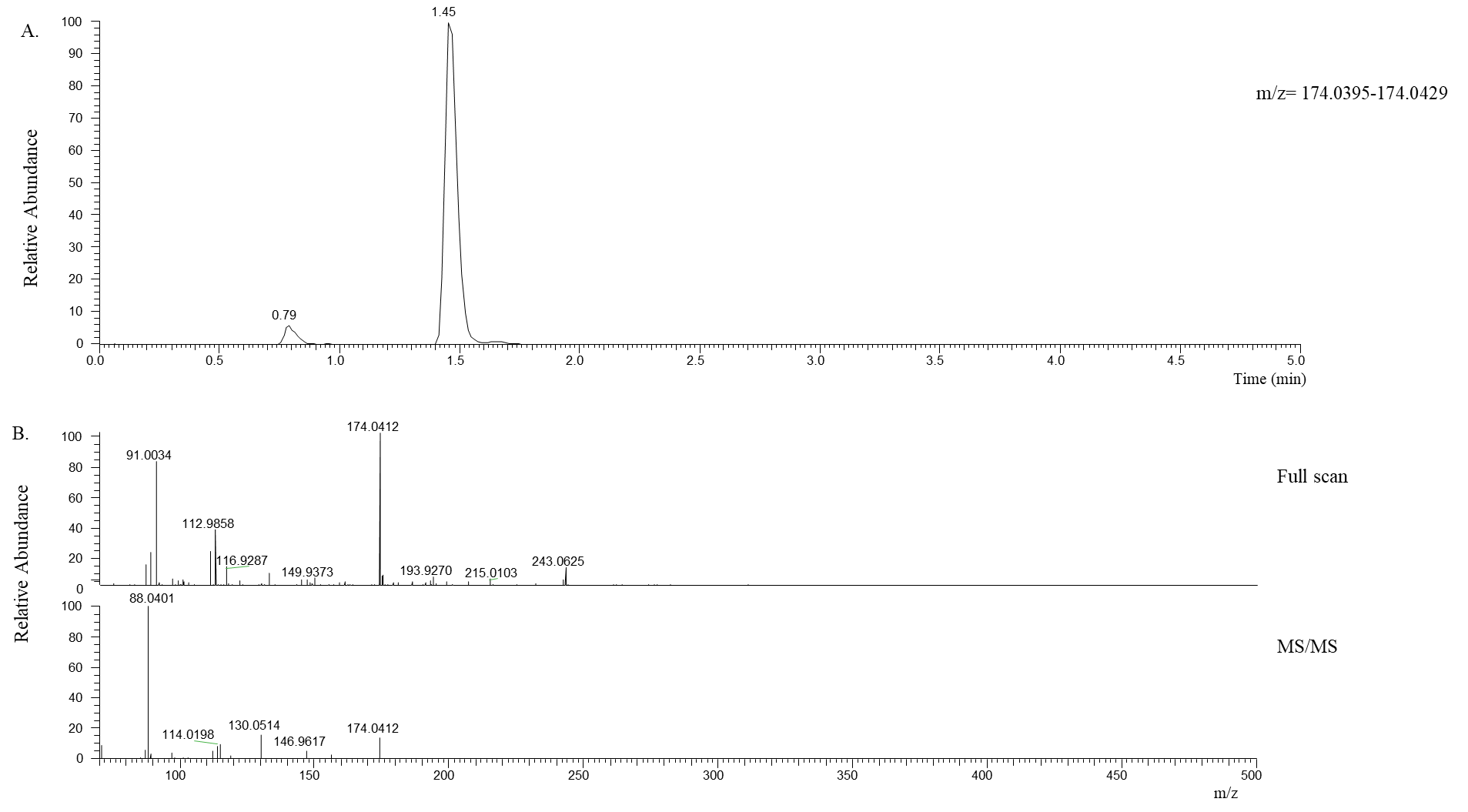

**Figure S22.**  Positive ionization for the identification of N-acetyl-L-aspartic acid.  Top, extracted ion chromatogram for m/z 174.0412 (5 ppm). Middle, full scan mass spectrum from retention time 1.45 min.  Bottom, tandem mass spectrum (MS/MS) for m/z 174.0412 (1.2 amu isolation window).

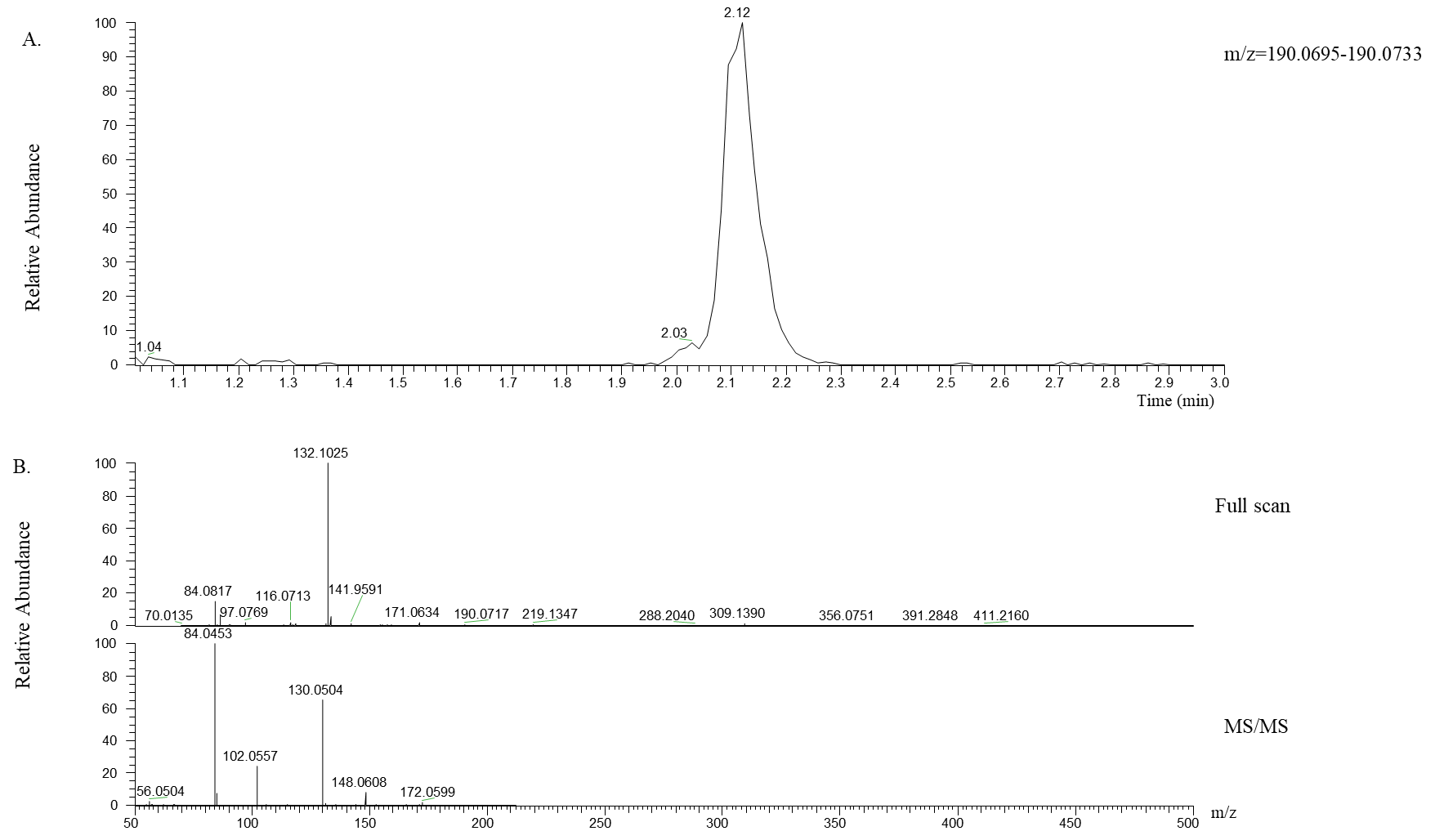

**Figure S23.**  Positive ionization for the identification of N-acetyl-L-glutamic acid.  Top, extracted ion chromatogram for m/z 190.0717 (5 ppm). Middle, full scan mass spectrum from retention time 2.12 min.  Bottom, tandem mass spectrum (MS/MS) for m/z 190.0712 (1.2 amu isolation window).

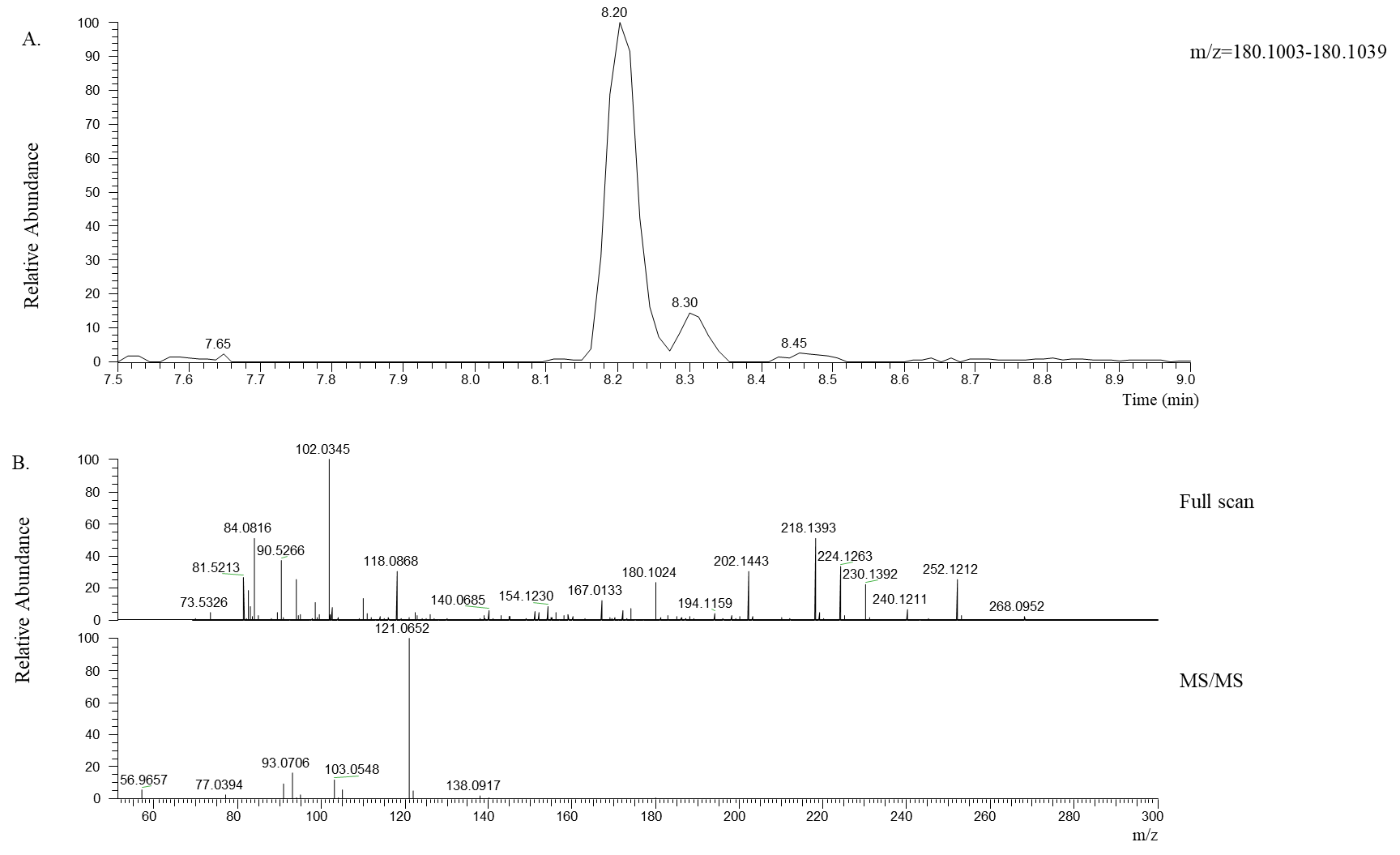

**Figure S24.**  Positive ionization for the identification of N-acetyltyramine.  Top, extracted ion chromatogram for m/z 180.1024 (5 ppm). Middle, full scan mass spectrum from retention time 8.20 min.  Bottom, tandem mass spectrum (MS/MS) for m/z 180.1024 (1.2 amu isolation window).

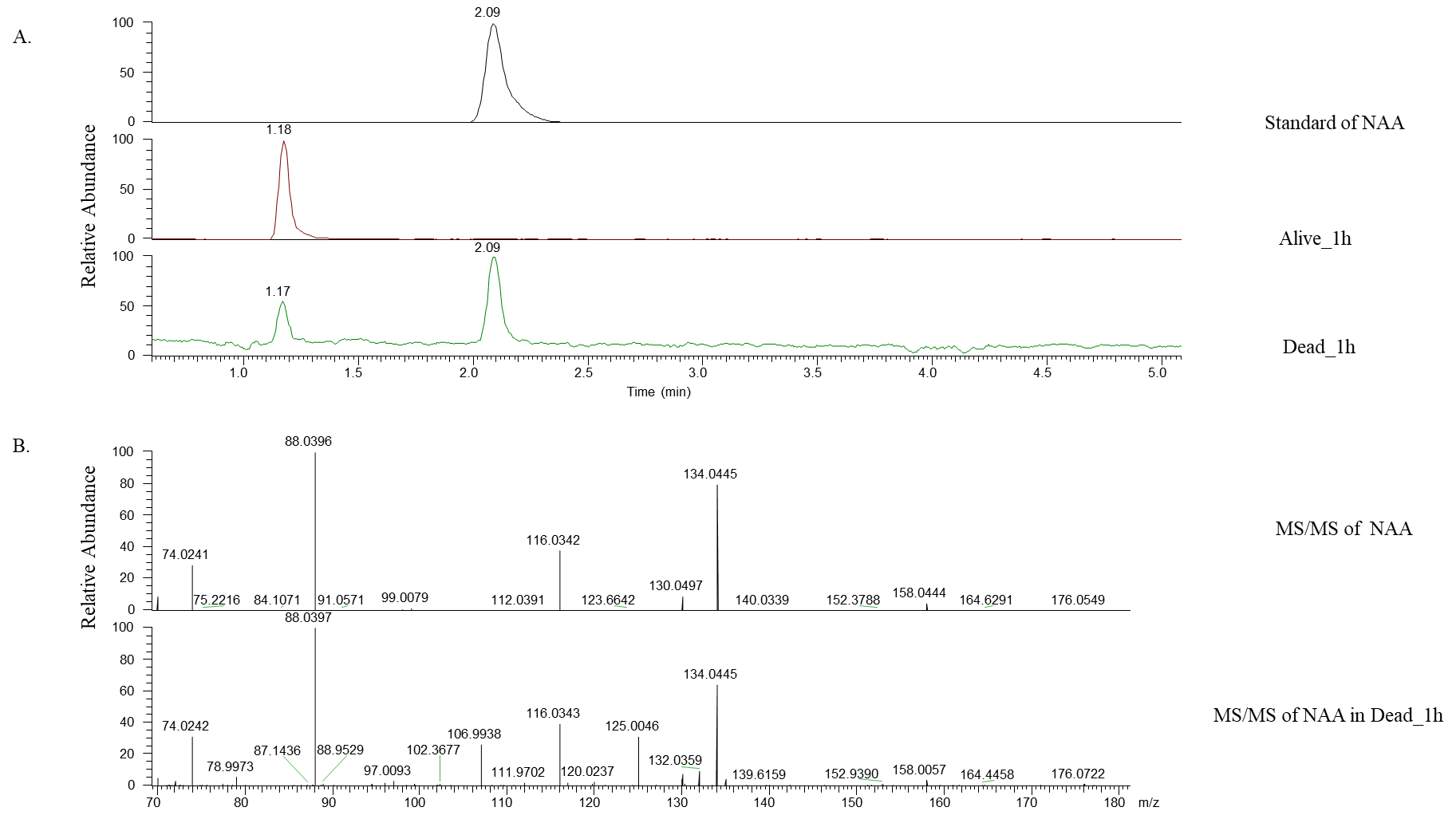

**Figure S25.**  Standard analysis of N-acetyl-L-aspartic acid (NAA)

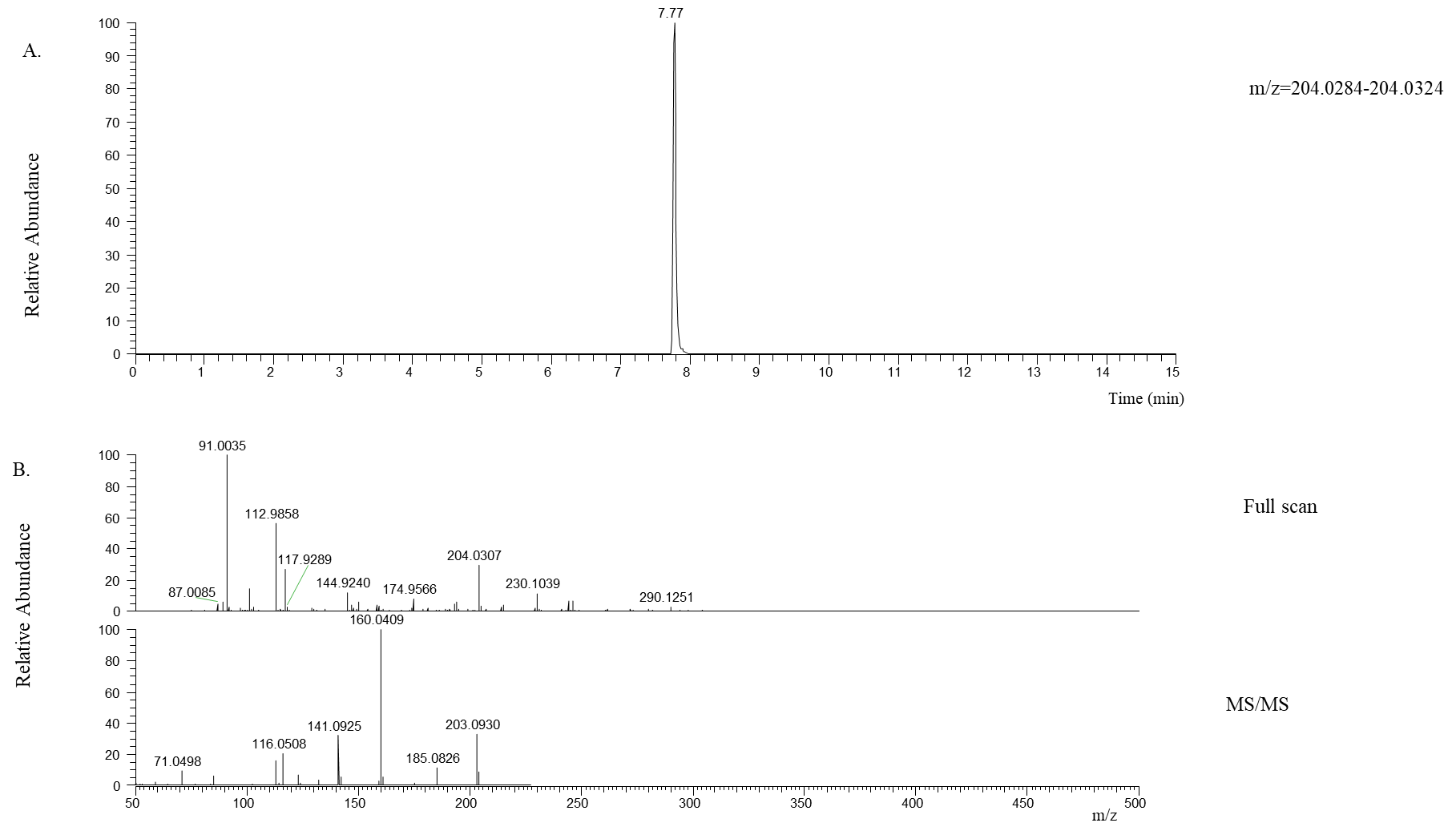

**Figure S26.**  Positive ionization for the identification of xanthurenic acid.  Top, extracted ion chromatogram for m/z 204.0307 (5 ppm). Middle, full scan mass spectrum from retention time 7.77 min.  Bottom, tandem mass spectrum (MS/MS) for m/z 204.0307 (1.2 amu isolation window).

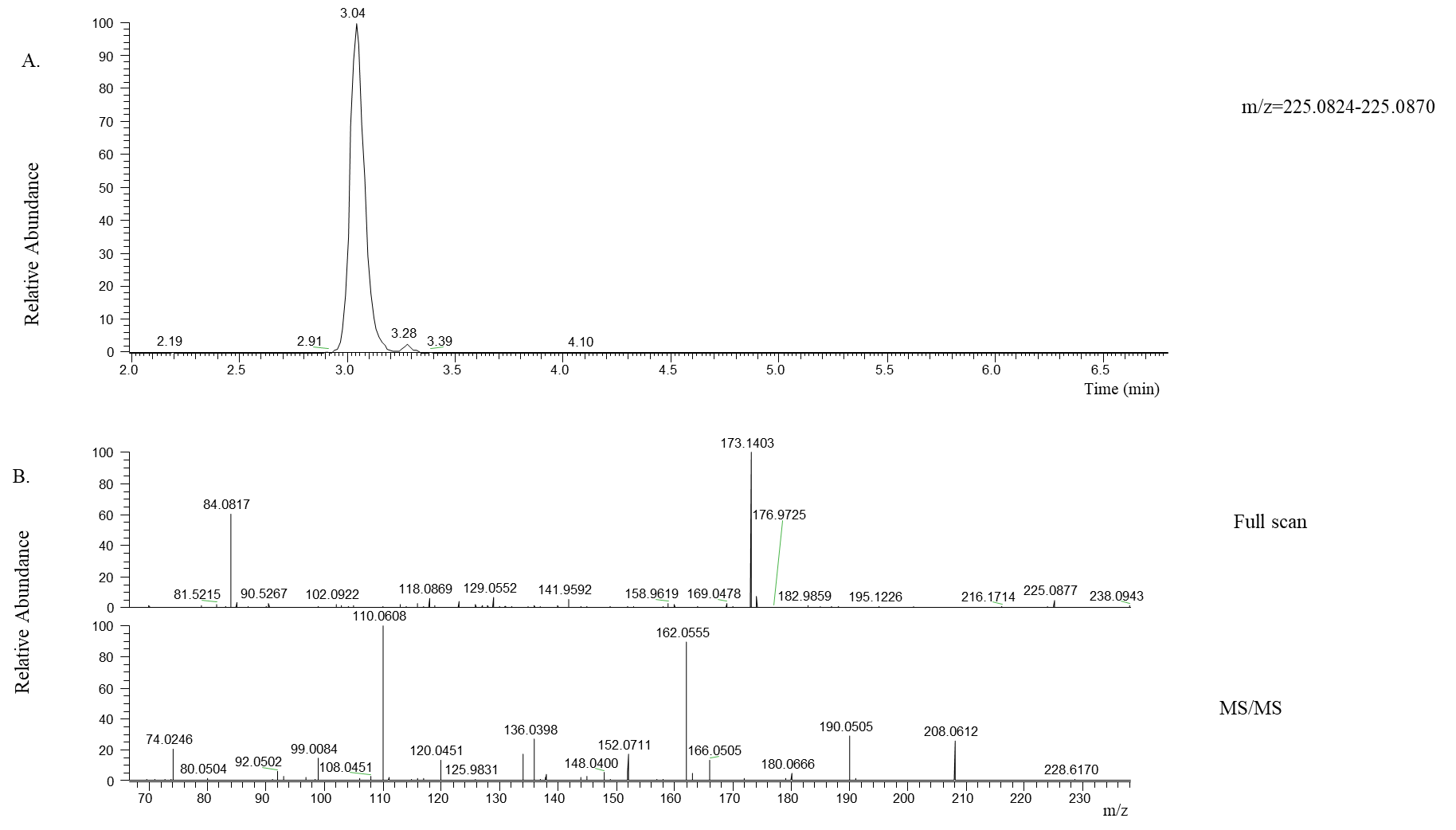

**Figure S27.**  Positive ionization for the identification of 3-hydroxykynurenine.  Top, extracted ion chromatogram for m/z 225.0877 (5 ppm). Middle, full scan mass spectrum from retention time 3.04 min.  Bottom, tandem mass spectrum (MS/MS) for m/z 225.0877 (1.2 amu isolation window).

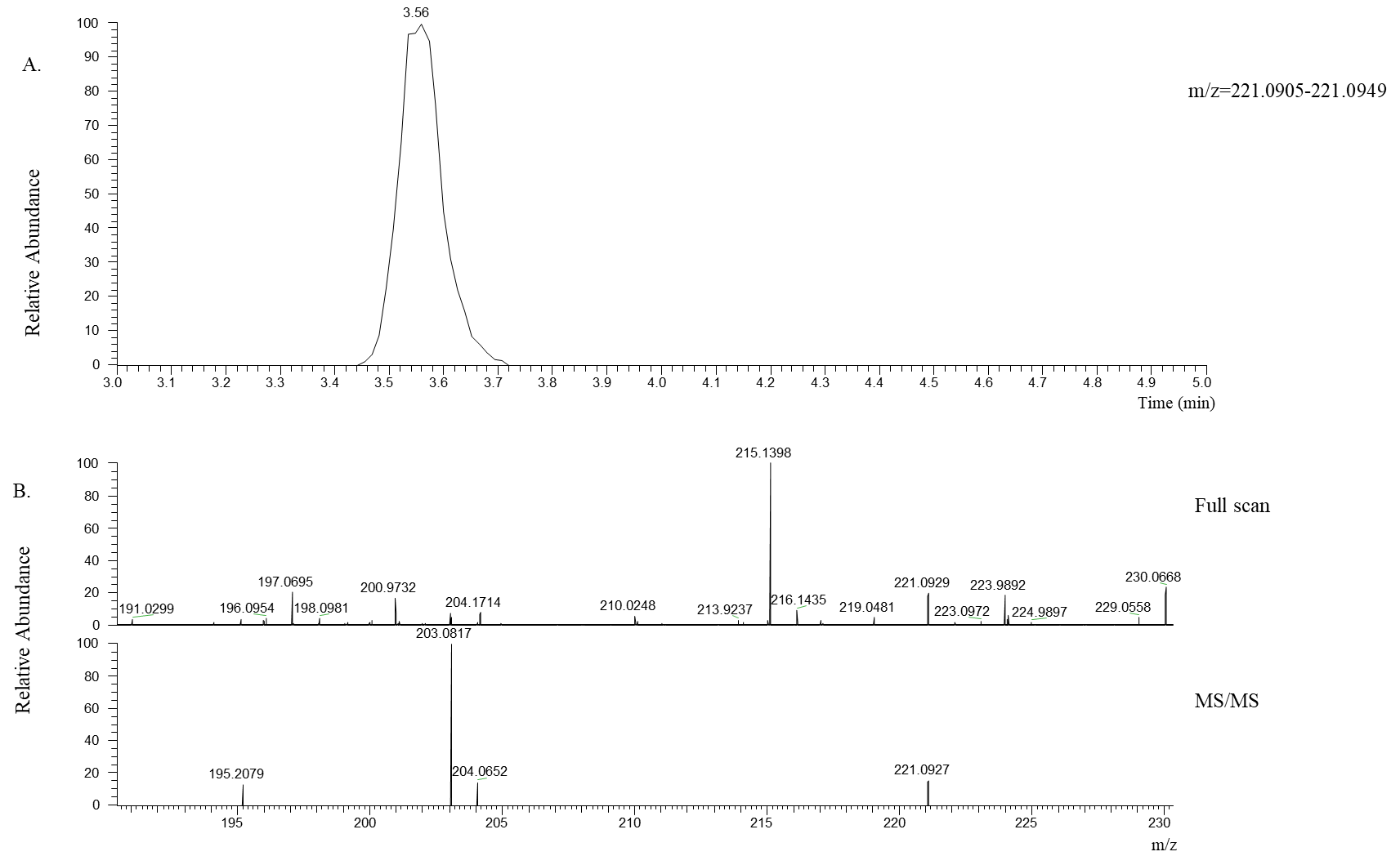

**Figure S28.**  Positive ionization for the identification of 5-hydroxyl-L-tryptophan.  Top, extracted ion chromatogram for m/z 221.0929 (5 ppm). Middle, full scan mass spectrum from retention time 3.56 min.  Bottom, tandem mass spectrum (MS/MS) for m/z 221.0929 (1.2 amu isolation window).

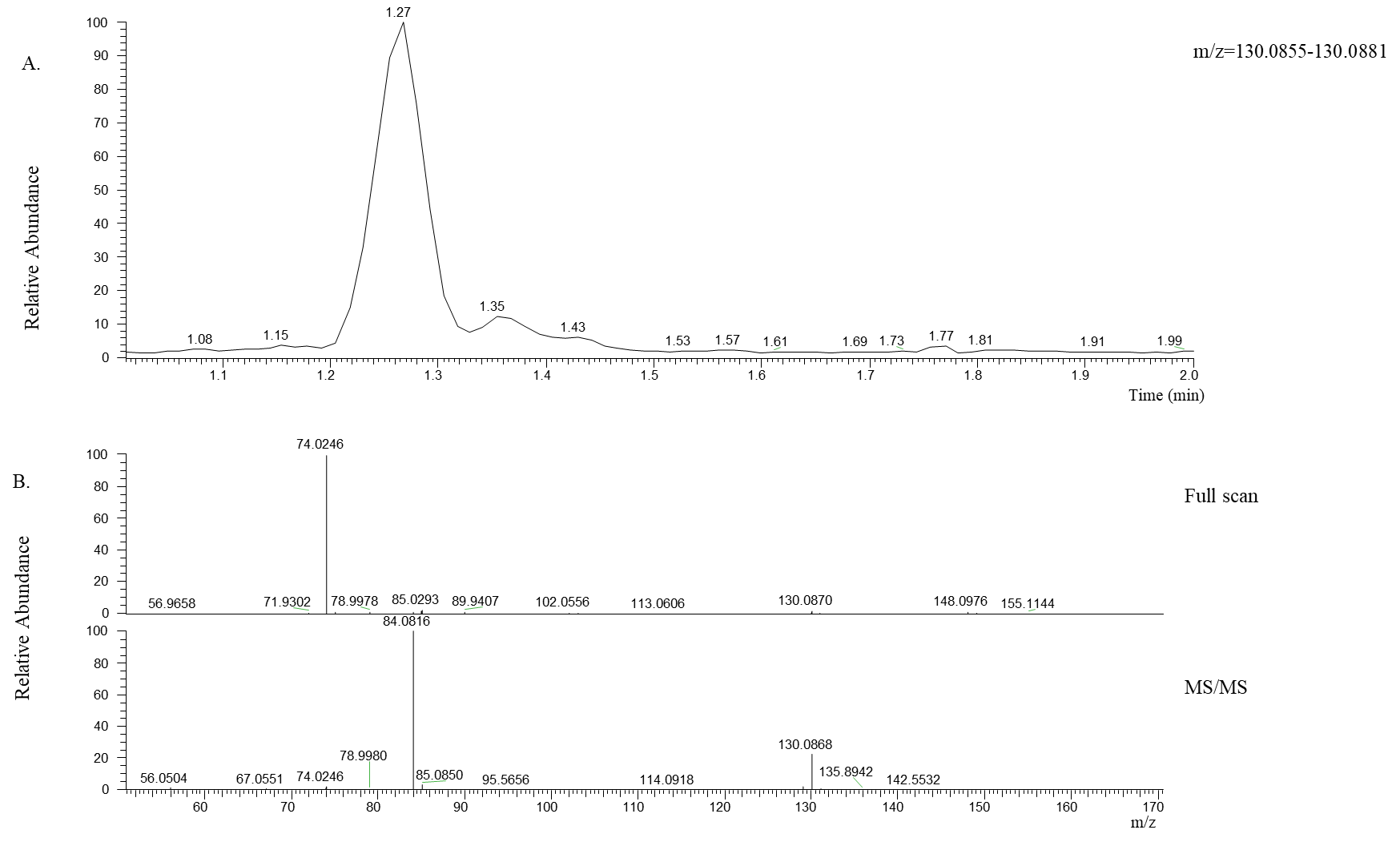

**Figure S29.**  Positive ionization for the identification of L-pipecolic acid.  Top, extracted ion chromatogram for m/z 130.0868 (5 ppm). Middle, full scan mass spectrum from retention time 1.27 min.  Bottom, tandem mass spectrum (MS/MS) for m/z 130.0868 (1.2 amu isolation window).

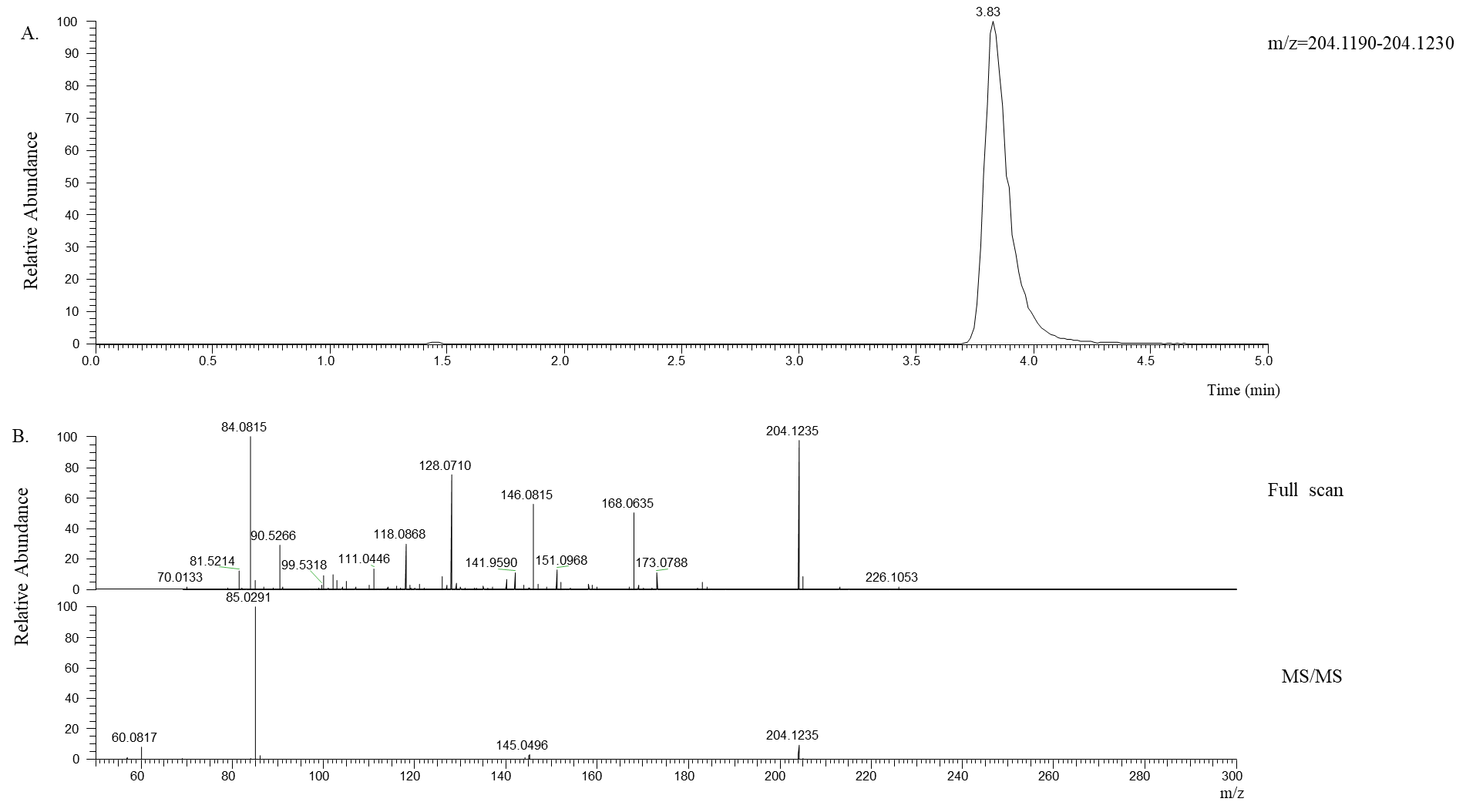

**Figure S30.**  Positive ionization for the identification of N-acetyl-L-carnitine.  Top, extracted ion chromatogram for m/z 204.1235 (5 ppm). Middle, full scan mass spectrum from retention time 3.83 min.  Bottom, tandem mass spectrum (MS/MS) for m/z 204.1235 (1.2 amu isolation window).

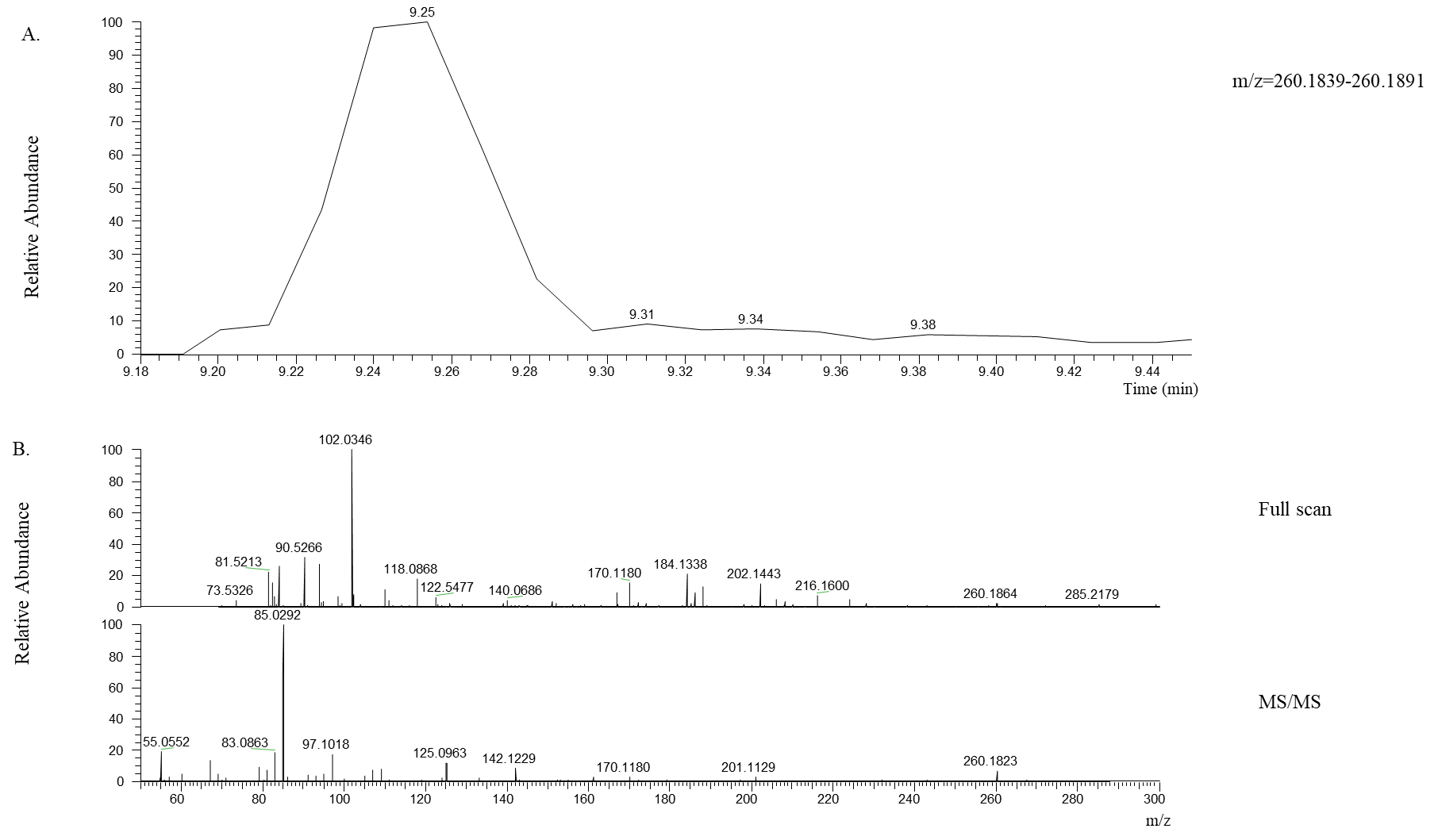

**Figure S31.**  Positive ionization for the identification of hexanoylcarnitine.  Top, extracted ion chromatogram for m/z 260.1864 (5 ppm). Middle, full scan mass spectrum from retention time 3.83 min.  Bottom, tandem mass spectrum (MS/MS) for m/z 260.1823 (1.2 amu isolation window).

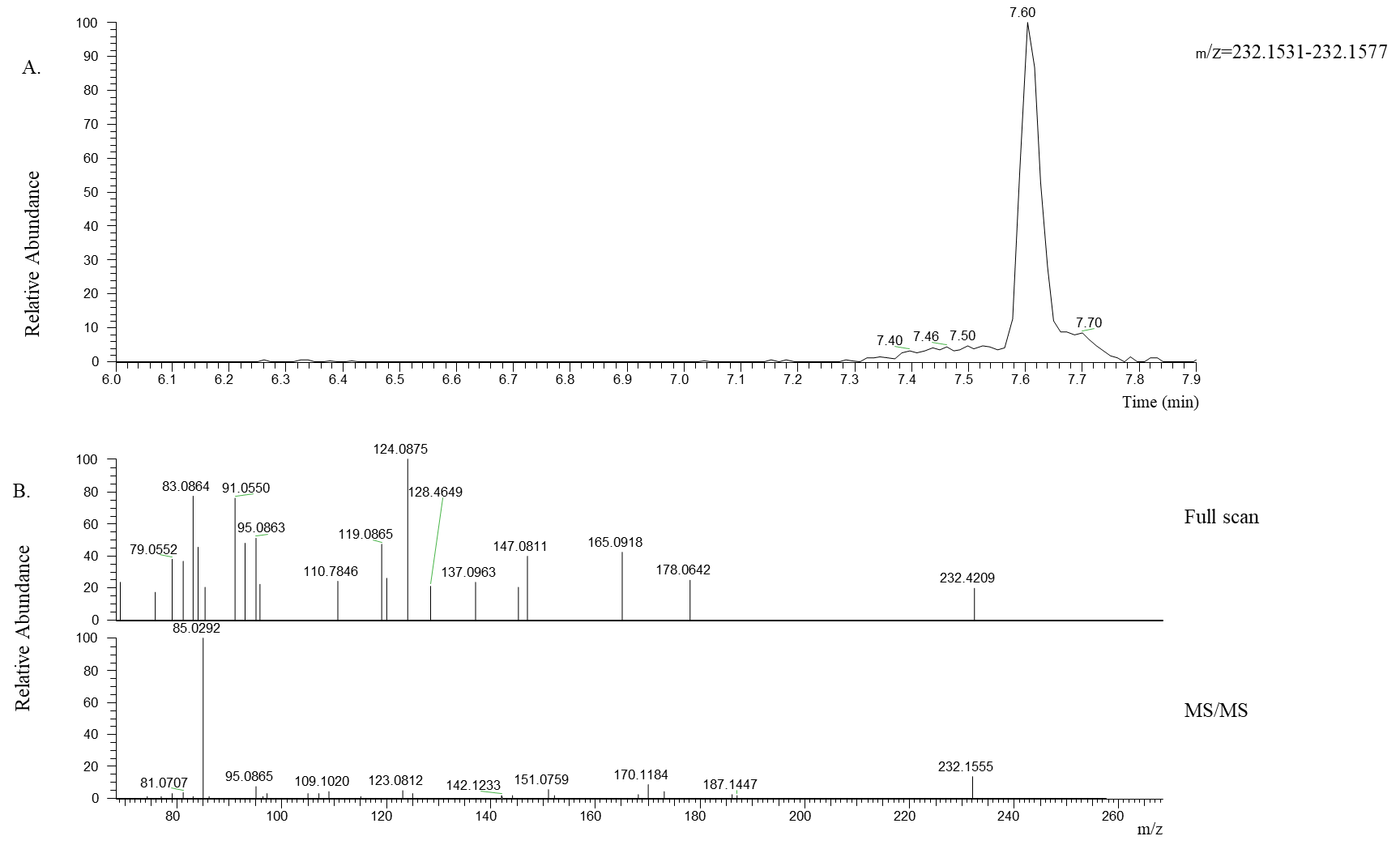

**Figure S32.**  Positive ionization for the identification of butanoylcarnitine.  Top, extracted ion chromatogram for m/z 232.4209 (5 ppm). Middle, full scan mass spectrum from retention time 7.60 min.  Bottom, tandem mass spectrum (MS/MS) for m/z 232.4209 (1.2 amu isolation window).

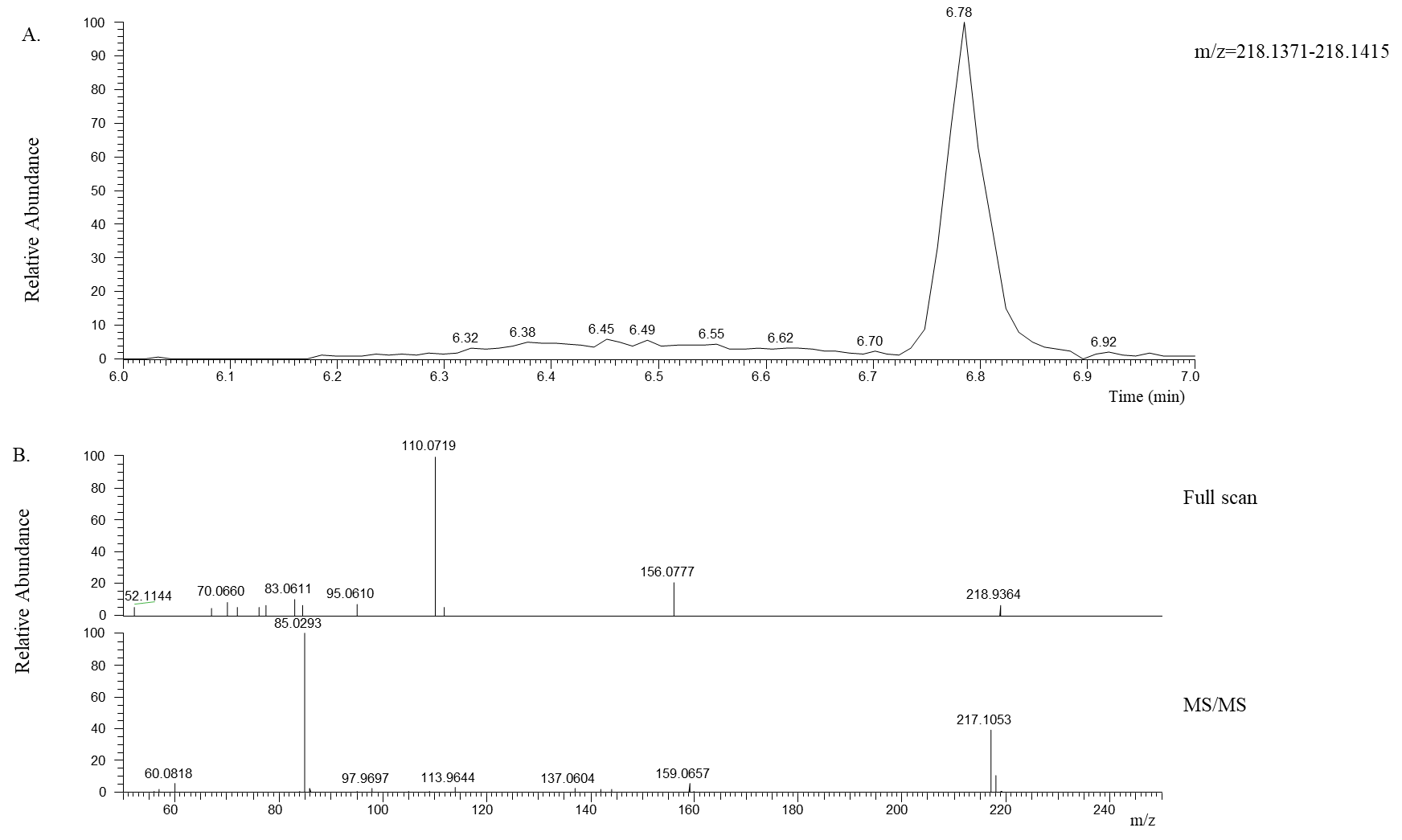

**Figure S33.**  Positive ionization for the identification of propanoylcarnitine.  Top, extracted ion chromatogram for m/z 218.9364 (5 ppm). Middle, full scan mass spectrum from retention time 6.78 min.  Bottom, tandem mass spectrum (MS/MS) for m/z 218.9364 (1.2 amu isolation window).
